# Airway applied IVT mRNA vaccine needs specific sequence design and high standard purification that removes devastating dsRNA contaminant

**DOI:** 10.1101/2024.09.22.614306

**Authors:** Jingjing Zhang, Chao Li, Yuheng Liu, Rui Liao, Dian He, Lifeng Xu, Tingting Chen, Qin Xiao, Mingxing Luo, Yang Chen, Yali Li, Huaxing Zhu, Joseph Rosenecker, Xiaoyan Ding, Shuchen Pei, Shan Guan

## Abstract

The development of next-generation mucosal mRNA vaccines is promising but extremely challenging. Major efforts have been focused on optimizing delivery systems, whereas it is still unknown whether the intrinsic quality of IVT mRNA significantly impacts the potency of airway inoculated mRNA vaccines. Here, we systematically demonstrate the mucosal mRNA vaccine requires a higher standard of purification and tailor-designed sequence to fulfil its potency compared to the parenteral route inoculated counterpart. We found double strand RNA (dsRNA) contaminants are prone to trigger innate immunoreaction in the airway that activates the mRNA degradation mechanism, thereby diminishing the mRNA expression and subsequent antigen-specific immune responses. To address these challenges, we developed a strategy that combines optimized untranslated regions (UTRs) screened from endogenous genes of pulmonary cells with affinity chromatography-based purification which removes almost all the dsRNA contaminants. The optimized mRNA administered via the airway route not only demonstrates superior protein expression (30-fold increase) and reduces inflammation in the lung, but also promotes robust immunity comprising significantly elevated systemic, cellular, and mucosal immune responses, which is in stark contrast to intramuscular injected counterpart that displays less pronounced benefits. Our findings offer new insight into the development of mucosal mRNA therapeutics from an overlooked but crucial perspective of optimizing mRNA components.

## Introduction

In just a few short years, Corona Virus Disease 2019 (COVID-19) vaccines based on in vitro transcribed messenger RNA (IVT mRNA) have been administered billions of times, saving millions of lives(*1*). The IVT mRNA technology has revolutionized the field of vaccine development due to its record-breaking speed of clinical translation as well as its flexible design to express any target protein(*2–4*). The approved mRNA vaccines have validated this platform and stimulate huge interests in developing next-generation mRNA vaccines, including mucosal mRNA vaccines that could elicit robust protections at the entry portal of a large number of notorious pathogens like Severe Acute Respiratory Syndrome Coronavirus 2 (SARS-CoV-2), influenza and *Mycobacterium tuberculosis*(*5, 6*). Almost all the approved vaccines are administered via injection, which triggers a strong systemic immune response but often leads to weak local mucosal immunity, potentially failing to prevent mucosal infections. In contrast, pulmonary vaccination not only generates robust mucosal immunity marked by secretory IgA (sIgA) and lung tissue resident memory T (Trm) cells but also induces strong systemic immune responses producing neutralizing antibodies in the serum, making it a promising strategy for preventing airborne pathogen colonization(*7–11*).

Despite their potential, developing mucosal mRNA vaccines is extremely challenging. Apart from a few viral vectors-based COVID-19 mucosal vaccines approved in China, Russia and India, only nine mucosal vaccines eight oral and one intranasal are approved worldwide for human use, all of which are based on either whole inactivated or live attenuated viral vectors(*12*). This discrepancy between parenteral and mucosal approaches primarily arises from the unique physiological structure and aggressive immune environment inherent to the respiratory mucosa(*13*). When applied to the airway, these barriers limit the effectiveness of currently available mRNA vaccines, which were originally designed for intramuscular injection. To address these challenges, the major efforts have been focused on optimizing mRNA delivery systems, such as improving the structure and compositions of lipid nanoparticle (LNP) or developing tailor-designed pulmonary delivery vehicles by leveraging biocompatible materials(*8, 14–19*). However, none of the previous studies, to the best of our knowledge, has revealed whether the properties of the mRNA molecule itself have a significant impact on the potency of mRNA vaccines inoculated via the respiratory tract. It is generally acknowledged that the properties of the mRNA molecule, such as its purity and sequence design, are able to impact parenteral mRNA vaccine potency(*20–23*). On the other hand, how these parameters affect the efficacy of mucosal mRNA vaccines is still unknown.

In this study, we systematically revealed that the mRNA quality, specifically sequence design and purity, is extremely crucial for the efficacy of airway-inoculated mRNA vaccine. Mucosal mRNA vaccines require tailor-designed sequences and a substantially higher standard of mRNA purification to fulfill their biological functions compared to intramuscularly injected counterparts. To optimize mRNA sequence design for mucosal vaccination, we focus on the 5 ′ untranslated region (UTR) and 3 ′ UTR structures flanking the coding sequence of IVT mRNA molecule as these regions play crucial roles in regulating mRNA stability and directly impact antigen expression levels(*21, 24–26*). To this end, we screened the 5’ UTR and 3’ UTR derived from highly mRNA-abundant genes in the pulmonary tissue using a bioinformatics analysis and found the expression of IVT mRNA in both cultured cells and the lung of mice were significantly enhanced when utilizing the UTRs of *TMSB10* gene. Additional modifications such as the insertion of functional elements and removal of microRNA (miRNA) binding sites on the UTRs, could further enhance mRNA translation efficiency in the lungs of mice to levels that are 10 times higher than counterpart adopting UTRs from the approved COVID-19 mRNA vaccine BNT-162b2. The sequence-optimized mRNA could mediate significantly improved levels of antigen-specific humoral, mucosal and cellular immune responses after the inhalation route (i.h.) of dosing compared to BNT-162b2 control, but failed to show substantial benefits within counterparts administered via the intramuscular route (i.m.), emphasizing the critical need for respiratory tract specific IVT mRNA sequence design.

We further confirmed a main culprit resulting in the distinct behaviors of intramuscular injected and respiratory route administered IVT mRNA is double-stranded RNA (dsRNA) impurities which are prone to be recognized by innate immune sensors such as 2’-5’oligoadenylate synthetase (OAS), toll-like receptor 3 (TLR3), retinoic acid-inducible gene I (RIG-I), and melanoma differentiation-associated protein 5 (MDA-5) as pathogen-associated molecular patterns (PAMPs) to trigger unwanted immune responses(*27–30*). The dsRNA contaminant containing IVT mRNA inoculated via i.h. route can significantly exacerbate these unintended side effects compared to i.m. injection, probably due to the high sensitivity of the respiratory tract to inflammation and its dense network of immune cells ready to respond to foreign substances(*31*). This heightened reactivity can lead to the activation of the mRNA degradation mechanism, which eliminates the IVT mRNA more rapidly, thereby resulting in a low level of the target antigen expression as well as a diminished antigen-specific immune response. Leveraging an affinity chromatography-based purification, we could successfully remove almost all of the dsRNA (>99.5%) from the IVT mRNA product, which in turn alleviates unwanted immune activation and improves mRNA transfection in the airway. Enhanced efficacy of purified mRNA in the scenario of respiratory route dosing is evidenced by a significant boost in target protein expression (30-fold), circulating IgG antibody (4-fold), CD8^+^IFN-γ^+^ T cells expansion (9-fold), and mucosal immunity as indicated by sIgA levels (4-fold) as well as CD8^+^ Trm cells (68-fold) compared to control groups, alongside reduced inflammation in lung and decreased proinflammatory cytokine secretion. In contrast, intramuscular administration of purified mRNA shows less pronounced benefits: target protein expression increased only 3-fold, IgG levels upregulated 2-fold, CD8^+^IFN-γ^+^ T cells improved 3-fold, with background level Trm cells and no detectable antigen-specific sIgA. Besides, we further confirmed the incorporation of dsRNA into purified IVT mRNA significantly impairs its in vivo expression, but with different impacts on i.m. (63.0% decrease) and i.h. (98.8% loss of function) routes.

Overall, this study provides clues to the reasons why developing mucosal mRNA vaccines is so challenging and highlights the detrimental repercussion of dsRNA in the respiratory tract. Our data underscores the potential of combining pulmonary-specific UTR design with affinity chromatography-based purification to enhance the immunogenicity of mucosal mRNA vaccines, offering valuable insights for advancing mRNA therapeutics targeting the respiratory tract(*17, 32, 33*).

## Results

### 1. Screening and modification of endogenous UTRs to enhance IVT mRNA translation

Given the vital roles of 5 ′ UTRs and 3 ′ UTRs within IVT mRNA sequence on the protein translation process, rational design on these structures holds great potential to improve the target antigen production from mucosal mRNA vaccines (Fig. 1A). Leveraging this biological principle, we conducted an analysis to rank gene mRNA abundance in respiratory tract related cell types ranging from bronchial epithelium to pulmonary dendritic cells (DCs), through utilizing RNA single cell type tissue cluster data that retrieved from the human protein atlas (HPA) (*34*). For each cell type, all genes were ranked using the normalized Transcripts Per Million (nTPM) parameter, and the top 20 genes were selected (Supplementary Table 1). It turns out that 10 of the selected genes, i.e. *ACTB, FTL, HSP90AA1, RPL18, TMSB10, RPS5, RPL6, RPS20, RPLP0* and *VIM*, were consistently expressed in all lung cell types (Fig. 1B and 1C). The 5′UTRs and 3′UTRs of these genes were selected and in vitro transcribed into a set of mRNA candidates encoding firefly luciferase (Fluc) (Supplementary Table 2 and Supplementary Table 3). mRNA translation was further conducted to evaluate the potential of different UTRs in respiratory tract-related cells, including 16HBE, DC2.4, RAW264.7 and A549 cells. Notably, UTRs of *TMSB10* demonstrated the highest mRNA translation efficacy across all cell types tested (Fig. 1D, Supplementary Fig. 1A and Supplementary Fig.1B). Based on these observations, UTRs from *TMSB10* were selected for artificial design to further optimize its potency.

**Fig. 1.**
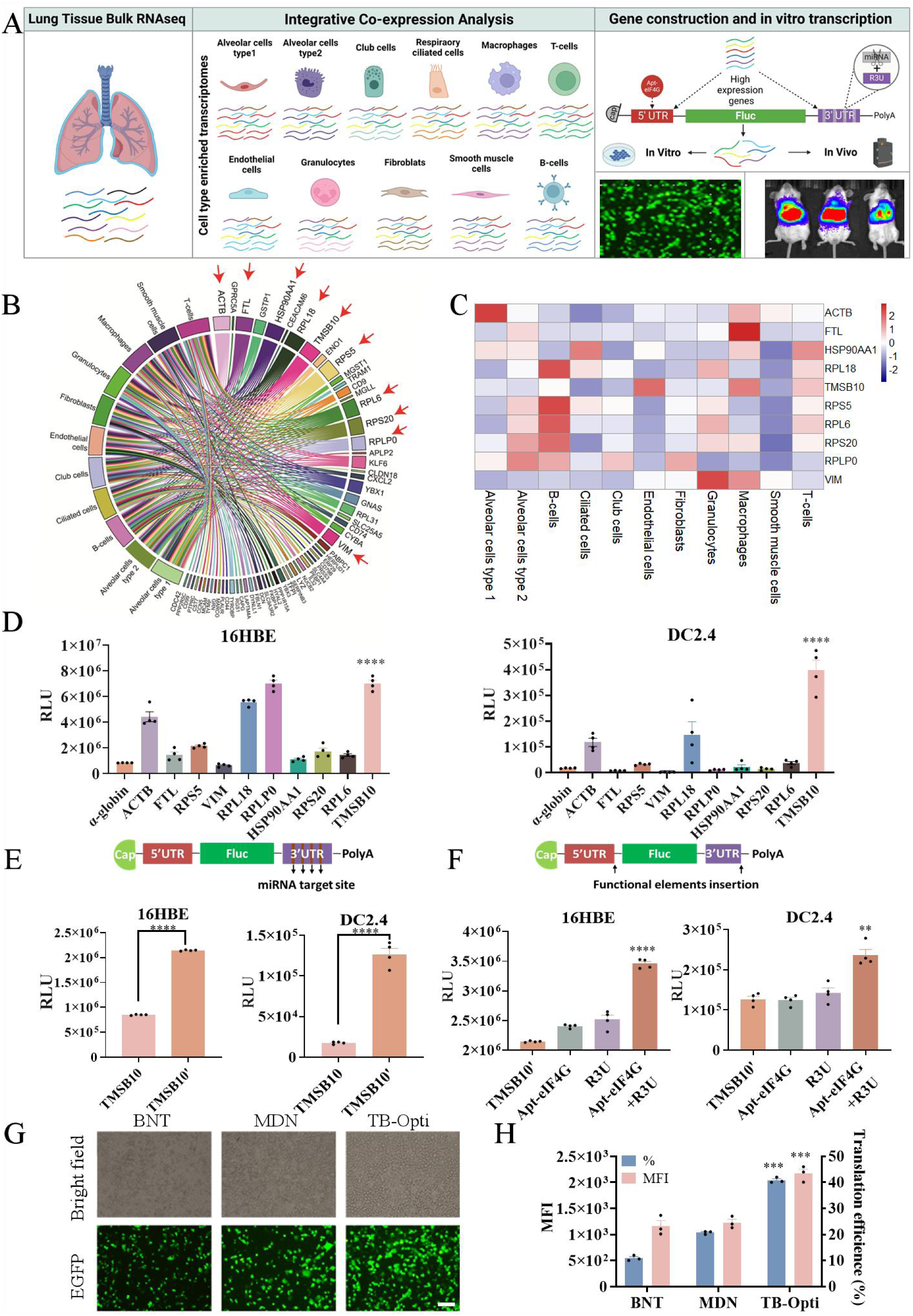
Screening and modification of IVT mRNA sequence for enhanced gene expression in respiratory tract-related cells. (**A**) Schematic illustration of the experimental workflow in screening and designing optimized mRNA sequence. This begins with an integrative co-expression analysis of lung tissue RNA sequencing data, identifying key genes expressed across various lung cell types, including epithelial cells, alveolar cells, endothelial cells, and immune cells. Selected 5’ UTR and 3’ UTR from top-ranked genes were incorporated into IVT mRNA constructs encoding firefly luciferase (mFluc), along with UTRs optimization by miRNA target site deletion and functional elements insertion. These constructs were then evaluated for transfection efficiency using both in vitro and in vivo models. (**B**) Chord plot illustrating the ranking of gene mRNA abundance across different respiratory tract-related cell types using data from the Human Protein Atlas (HPA). The top 20 genes from each cell type were selected for further analysis. The red arrows refer to genes that were consistently expressed in all lung cell types. (**C**) Expression heatmap of selected genes across lung cell types, with consistent expression patterns observed for the final set of 10 genes, including *TMSB10*. (**D**) The IVT mRNA constructs were designed with different 5’ UTRs and 3’ UTRs derived from selected genes and transfected into 16HBE and DC2.4 cells to assess their impact on mRNA transfection efficiency. (**E**) Schematic illustration of a truncated version of TMSB10 3’ UTR (TMSB10’) designed to remove potential miRNA target sites and evaluation via mFluc transfection in 16HBE and DC2.4 cells. (**F**) Schematic illustration of functional motifs integrated into TMSB10’ UTRs, including Apt-eIF4G in the 5’ UTR and R3U in the 3’ UTR. The mFluc with modified UTRs were tested in 16HBE and DC2.4 cells. (**G-H**) Fluorescence microscopy (G) and flow cytometry analysis (H) evaluating the expression levels of mRNA encoding EGFP (mEGFP) designed with UTRs of TB-Opti, BNT and MDN. TB-Opti: IVT mRNA using the above mentioned best-performing UTRs of TMSB10-optimized design, BNT: IVT mRNA with UTRs from the approved COVID-19 mRNA vaccine BNT-162b2, MDN: IVT mRNA with UTRs from the approved COVID-19 mRNA vaccine mRNA-1273 (more details can be found in Supplementary Table 2 and Supplementary Table 3). Statistical significance was assessed by two-tailed unpaired *t*-test and one-way ANOVA with Dunnett’s post-hoc test (\*\**P* < 0.01; \*\*\**P* < 0.001; \*\*\*\**P* < 0.0001). 16HBE: human bronchial epithelial cells; DC2.4: mouse bone marrow-derived dendritic cells; RAW264.7: mouse mononuclear macrophage leukemia cells; A549: human non-small cell lung cancer cells; RLU: relative light unit.

MicroRNA (miRNA) was considered to mainly target the 3’UTR of mRNA for negative gene regulations(*35*). Therefore, the possible miRNA target sites within 3’UTR of TMSB10, which were predicted using an online database-miRDB, were removed to create a variant termed TMSB10’. This modification led to a significant increase in protein expression approximately 1.5-fold in 16HBE cells and 6.2-fold in DC2.4 cells compared to the original TMSB10 construct (Fig. 1E). To further enhance mRNA translation, we selected two classic motifs for functional insertion to improve the potency of TMSB10’. The first functional motif, Apt-eIF4G, is an eIF4G recruiting aptamer that has been shown to increase mRNA translation when added to the 5’UTR(*36*). Another candidate motif is R3U (the repeated sequence element 3 and U-rich element), which originates from a Sindbis Virus RNA region responsible for RNA-binding proteins (RBP) binding to induce the potent stabilization of the viral RNA genome(*37*). These modifications, both individually and combined (Apt-eIF4G+R3U), significantly increased the levels of mRNA translation, with the combined modification showing the most substantial improvement (Fig. 1F). The optimized UTRs originating from TMSB10 with the above-mentioned modifications were finally adopted to synthesize IVT mRNA encoding various proteins of interest (Supplementary Fig. 1C) for all following investigations and termed as TMSB10-optimized (TB-Opti). To further confirm the superiority of TB-Opti, we adopted enhanced green fluorescent protein (EGFP) as a reporter and compared the TB-Opti with the UTRs design of the approved COVID-19 mRNA vaccines from both BioNtech BNT-162b2 (BNT) and Moderna mRNA-1273 (MDN). The fluorescence microscopy indicates more cells successfully translated in the TB-Opti group compared with BNT and MDN counterparts (Fig. 1G, Supplementary Fig. 8). Quantitative assessment via flow cytometry analysis confirmed a similar trend in which the TB-Opti group achieved a substantially higher ratio of EGFP-positive cells and significantly enhanced levels of mean fluorescence intensity (MFI) than BNT and MDN counterparts (Fig. 1H).

### 2. The impact of UTR sequence optimization on immune responses of mRNA vaccines inoculated via different routes

To assess the effectiveness of the TB-Opti sequence design, IVT mRNA encoding mFluc was encapsulated into LNPs for in vivo applications. When administered via i.h. route, mFluc using TB-Opti sequence design produced substantially elevated levels of bioluminescence in both live mice and excised lungs compared to BNT counterpart (Fig. 2A). Although moderate increases of transfection could be detected in TB-Opti mFluc-treated mice after intramuscular route (i.m.) dosing, there is no significant difference between TB-Opti mFluc treated group and counterpart adopting BNT design neither in live animals nor in excised organs (Fig. 2B). Afterwards, we synthesized IVT mRNA encoding the receptor binding domain (RBD) of the SARS-CoV-2 Omicron variant using TB-Opti design (mRBD, details can be found in Supplementary Table 2, Supplementary Table 3 and Supplementary Table 4) and immunized mice via different routes using an immunization scheme outlined in Fig. 2C. The serum samples collected from mice vaccinated by TB-Opti via i.h. route demonstrated a substantial increase of RBD-specific IgG antibody compared to the BNT counterpart (Fig. 2D), while no significant difference was observed in serum IgG levels between the TB-Opti and BNT groups when administered via i.m. route (Fig. 2E). Additionally, only the i.h. route inoculated mice displayed observable mucosal immune response, which is characterized by RBD-specific sIgA antibody in bronchoalveolar lavage fluid (BALF) and nasal lavage fluid (NLF) samples, with the TB-Opti groups outperforming BNT counterpart in both samples (Fig. 2F and 2G). Subsequently, spleens and lungs of vaccinated mice were collected to evaluate cellular immune response via the ELISpot assay, focusing on interferon-γ (IFN-γ), interleukin-4 (IL-4), and interleukin-17A (IL-17A) secreting cells. The levels of IFN-γ and IL-17A secretion in pulmonary lymphocytes from mice immunized with TB-Opti via i.h. route were significantly higher than the BNT counterpart (Fig. 2H). However, the elevated trend of IFN-γ secretion between TB-Opti and BNT was not observed in the groups receiving i.m. injections (Fig. 2I). These findings suggest that the TB-Opti prominently enhanced the immunogenicity of mRNA vaccines when applied via respiratory administration, implying specific mRNA sequence design is necessary for airway-inoculated mRNA vaccines. Based on these results, we selected the TB-Opti sequence design for all IVT mRNA in following investigations.

**Fig. 2.**
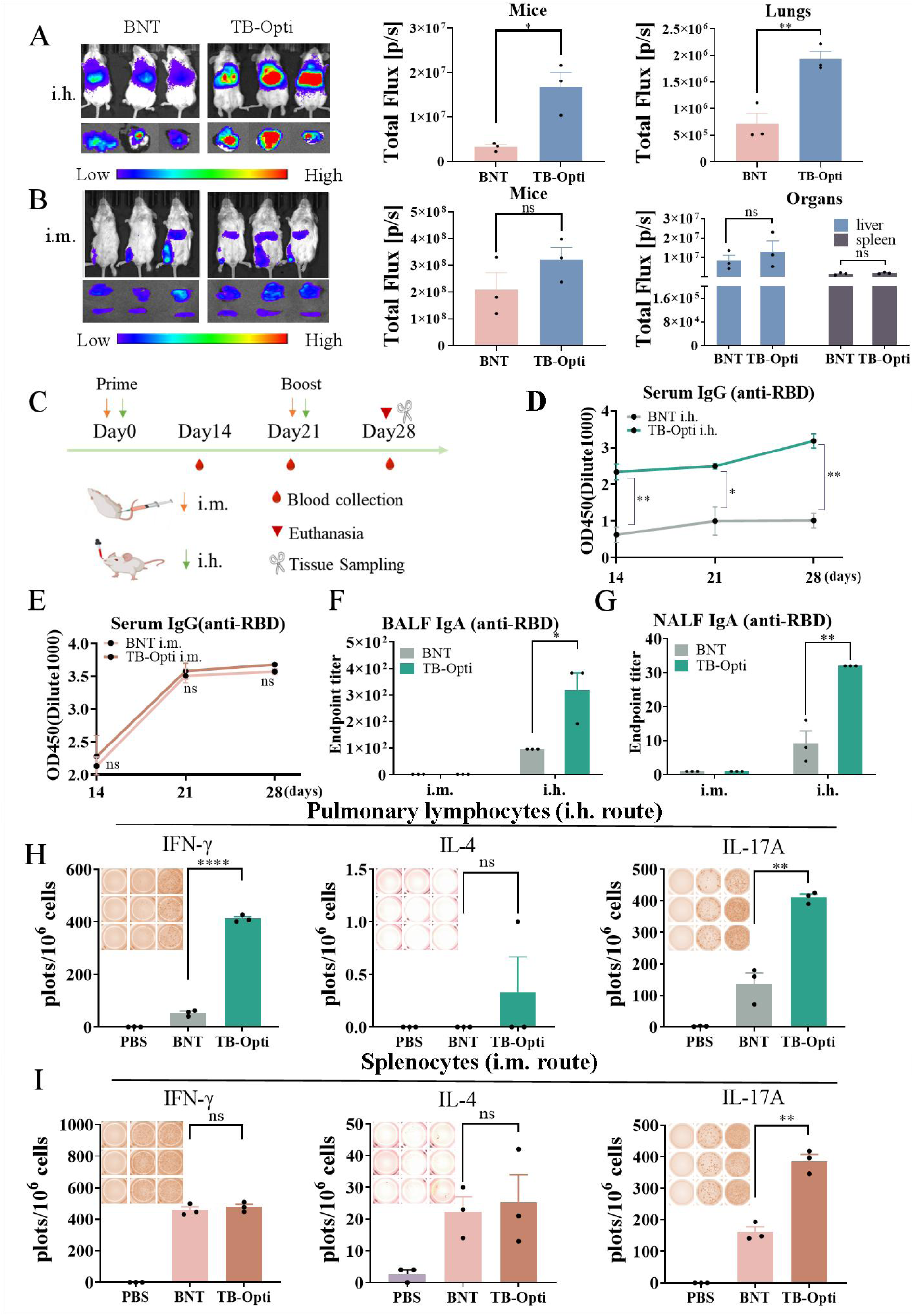
In vivo evaluations of IVT mRNA with TB-Opti sequence design. (**A-B**) Representative bioluminescence imaging of mRNA expression induced by mFluc using TB-Opti or BNT sequence design in mice and excised organs (left panel) via i.h. (A) or i.m. (B) route of dosing, with corresponding quantitative fluorescence intensities (middle and right panel), n = 3 biologically independent animals. (**C**) Schematic illustration of immunization, sampling and experimental procedure. (**D-E**) Optical density values (OD_450_) of RBD-specific IgG in serum samples from i.h. route (D) or i.m. route (E) vaccinated mice on day 14, day 21 and day 28. (**F-G**) Endpoint titers of the RBD-specific sIgA in BALF (F) and NLF (G) samples collected from mRBD with BNT or TB-Opti sequence design vaccinated mice on day 28. (**H-I**) ELISpot assay of IFN-γ, IL-4 and IL-17A spot-forming cells in (H) pulmonary lymphocytes (i.h. route) and (I) splenocytes (i.m. route) of vaccinated mice after re-stimulation with peptide pools of 14-mer overlapping peptides spanning the SARS-CoV-2 RBD region (details can be found in Supplementary Table 6). Statistical significance was calculated by two-tailed unpaired *t*-test (ns: not significant; \**P* < 0.05; \*\**P* < 0.01; \*\*\*\**P* < 0.0001). BNT represents IVT mRNA with UTRs from the approved COVID-19 mRNA vaccine BNT-162b2. TB-Opti represents mRNA with the best-performing UTRs developed in Fig. 1 (more details can be found in Supplementary Table 2 and Supplementary Table 3). i.h.: inhalation, i.m.: intramuscular injection.

### 3. Improving the quality of IVT mRNA via affinity chromatography-based purification

Previous publications suggest the purification of IVT mRNA may influence the antigen expression after the parenteral routes of dosing, necessitating the removal of contaminants generated during in vitro transcription(*29, 38*). As a result, we developed an Oligo-dT18 chromatographic column-based purification strategy (Fig. 3A). In this study, the “purified” means higher standard purification of IVT mRNA by affinity chromatography purification after LiCl precipitation, and the “unpurified” refers to IVT mRNA that is only purified by LiCl precipitation method. The details of the affinity chromatography purification are outlined in Fig. 3A and Supplementary Table 5. The oligo-dT18 column could successfully capture target IVT mRNA through interactions between their polyadenylated tails and thymine residues in a high-salt environment of mobile phase A (Fig. 3B). It was followed by efficient removal of impurities, including unincorporated nucleobases, RNAs with unwanted structures (including dsRNA), and proteins(*29*), via a mild phosphate buffer system (mobile phase B). The purified IVT mRNAs were subsequently eluted with nuclease-free water (mobile phase C) and concentrated at room temperature (RT) under vacuum conditions (Supplementary Fig. 3A). After purification, we utilized mobile phase D to clean the column, ensuring no residual impurities remained. The integrity of mRNAs with different lengths was confirmed through agarose gel electrophoresis, as evidenced by the presence of sharp bands shown in the purified group (Fig. 3C), implying different types of IVT mRNA with different lengths could be successfully purified by our strategy. Post-affinity chromatography, dsRNA depletion was validated via dot blot assay using dsRNA-specific antibodies, where the purified mRNA group (P) exhibited notably lower dsRNA levels compared to the unpurified counterpart (U) with almost all of dsRNA (>99.5%) successfully removed from purified mRNA group (P) (Fig. 3D).

**Fig. 3.**
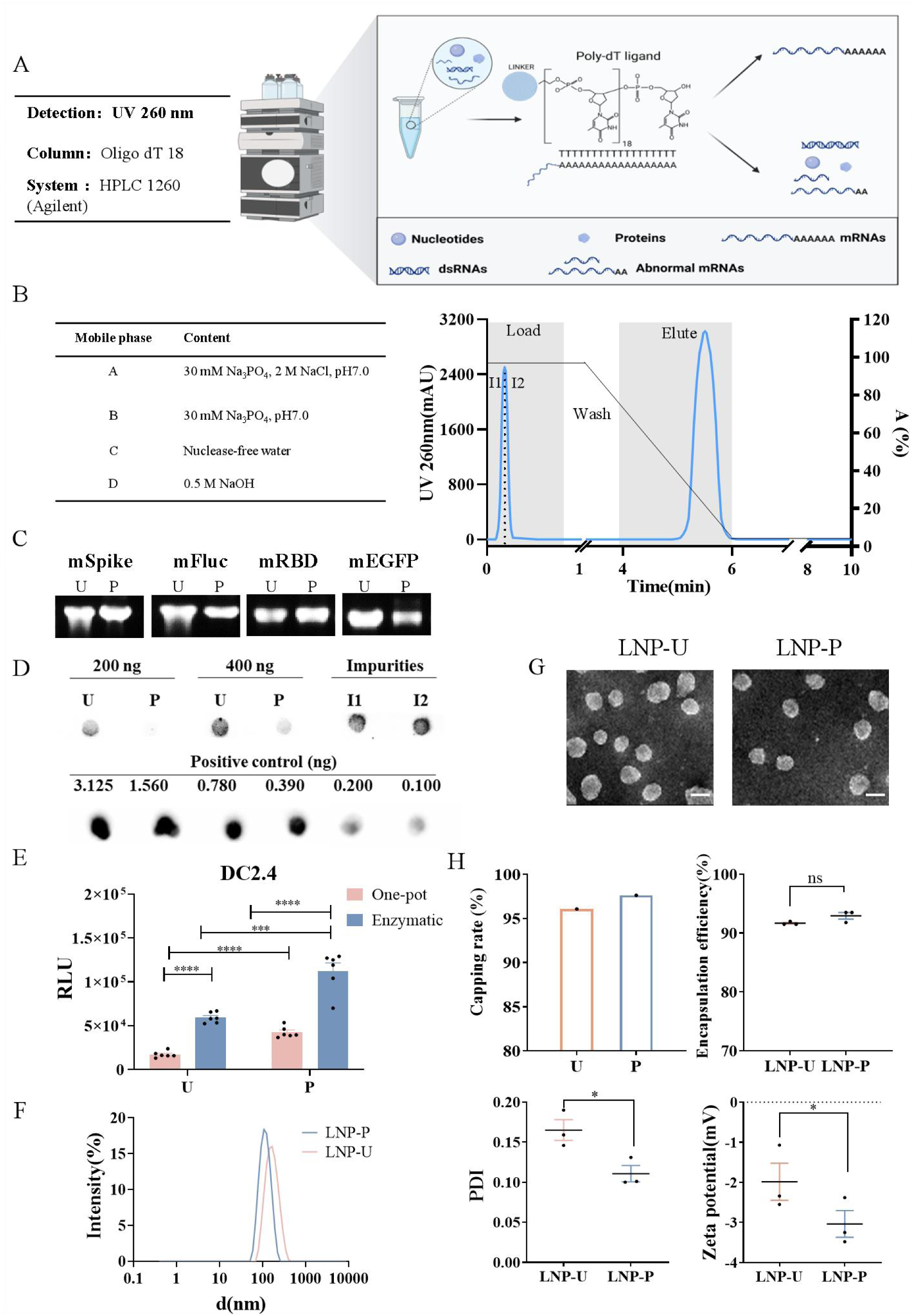
IVT mRNA purification with Oligo-dT-based affinity chromatography. (**A**) Schematic diagram of Oligo-dT chromatogram based IVT mRNA purification process with device parameters. (**B**) Detailed illustration of parameters applied in the Oligo-dT-based affinity chromatography purification. The IVT mRNA sample pre-purified by LiCl is loaded in the loading step. Unbound species such as nucleotides, plasmids, capping reagents, enzymes, and non-polyadenylated RNAs were washed out in the wash step. Polyadenylated mRNA elutes in the elution step. (**C**) Electrophoresis of unpurified (U) and purified (P) IVT mRNAs with different coding sequences and lengths. (**D**) The dot blot analysis on the purity of unpurified (U) and purified (P) IVT mRNA (200 ng or 400 ng). I1 and I2 represent different batches of impurities not bound to the column during the loading step in B. The positive control refers to the dsRNA standard (0.100-3.125 ng) prepared by the Novoprotein Scientific.Inc. (**E**) The transfection efficiency of IVT mRNA before (U) and after (P) purification in DC2.4 cells prepared by different capping methods. The “one-pot” method integrates IVT mRNA synthesis and capping within a single reaction, streamlining the process. In contrast, the “enzymatic” method involves first IVT mRNA transcription then capping in another separate step using a capping enzyme, which allows for more controlled and precise cap addition. (**F**) Dynamic light scattering measurement (hydrodynamic size, PDI and zeta potential) of LNP formulations loaded with unpurified IVT mRNA (LNP-U) or purified IVT mRNA (LNP-P). (**G**) Transmission electron microscopy images of LNP-U and LNP-P. Scale bar, 100 nm. (**H**) Capping rate of unpurified IVT mRNA (U) and purified IVT mRNA (P) (left panel), and encapsulation efficiency of LNP-U and LNP-P (right panel). Statistical significance was assessed by two-tailed unpaired *t*-test and one-way ANOVA with Dunnett’s post-hoc test (ns: not significant; \**P* < 0.05; \*\*\**P* < 0.001; \*\*\*\**P* < 0.0001).

Intriguingly, affinity chromatography-based purification strategy showed significant bias in the purification of IVT mRNAs prepared by “one-pot capping” (simultaneous transcription and capping) versus “enzymatic capping” (first transcription then capping in another separate step), with a preference for enzymatically capped mRNAs as proved in Fig. 3E. Based on this observation, all the IVT mRNAs applied in following investigations were prepared via enzymatic capping approach. We further evaluated the properties of purified IVT mRNAs versus unpurified counterparts after encapsulation in LNP, including their particle size, zeta potential, and polydispersity index (PDI). The LNP encapsulating the purified mRNA (LNP-P) exhibited better uniformity and smaller sizes than those from counterpart with unpurified mRNA (LNP-U); both groups predominantly exhibited as negatively charged nanoparticles (Fig. 3F). Additionally, the purification process had minimal impact on the morphology of the nanoparticles (Fig. 3G), the encapsulation efficiency of LNP formulations and the IVT mRNA capping rates (Fig. 3H), implying the advantages of this purification approach in maintaining the structural and functional integrity of IVT mRNA within LNP. All these results indicate that we successfully developed an affinity chromatography-based IVT mRNA purification method by which unwanted impurities, including dsRNA, could be efficiently removed in order to obtain high-quality IVT mRNA.

### 4. Purified IVT mRNA improvs translation efficiency and safety profiles in cultured cells

We loaded unpurified mFluc (mFluc-U) or purified mFluc (mFluc-P) into two types of commercially available in vitro transfection reagents, i.e. a cationic lipid-based Lipo8000 and a cationic polymer-based polyethyleneimine (PEI), to transfect various types of cultured cells. Compared to mFluc-U group, mFluc-P displayed significantly increased Fluc expression across all types of tested cells, and Lipo8000 based carriers could better mediate both mFluc-U and mFluc-P in transfecting the cultured cells than PEI based counterparts (Fig. 4A). Remarkably, the enhancement of mFluc-P could persist in 16HBE cells at least 96 h post-transfection, resulting in a significant overall improvement in Fluc expression (Fig. 4B). Western blot analysis further revealed significantly enhanced RBD expression of purified mRBD (mRBD-P) compared with unpurified mRBD (mRBD-U) in DC2.4 cells (Fig. 4C). The fluorescence microscopy demonstrated the number of cells expressing EGFP significantly increased in the purified group (mEGFP-P) compared to the unpurified counterpart (mEGFP-U) both in 16HBE and DC2.4 cells (Fig. 4D, Supplementary Fig. 3B). Quantitative assessment via flow cytometry analysis confirmed that the purified mEGFP-P successfully transfected approximately 50% 16HBE cells at 24 h post-transfection, concurrent with significantly increased MFI (Fig. 4E). In contrast, less than 36% 16HBE cells displayed significantly weaker fluorescence signals in mEGFP-U group (Fig. 4E). We further evaluated the cytotoxicity of purified IVT mRNA using the cell counting kit 8 (CCK8) assay, which indicated negligible impact of the purification process on the viability of 16HBE and DC2.4 cells (Fig. 4F). Additionally, the mFluc-P exhibited significantly reduced levels of inflammatory cytokines including interferon-α (IFN-α), interferon-β (IFN-β) and interleukin-6 (IL-6) (IFN-α, *P* < 0.01; IFN-β, *P* < 0.001; IL-6, *P* < 0.05) (Fig. 4G), indicating that the purification process effectively mitigates inflammatory stimulating profiles of IVT mRNA.

**Fig. 4.**
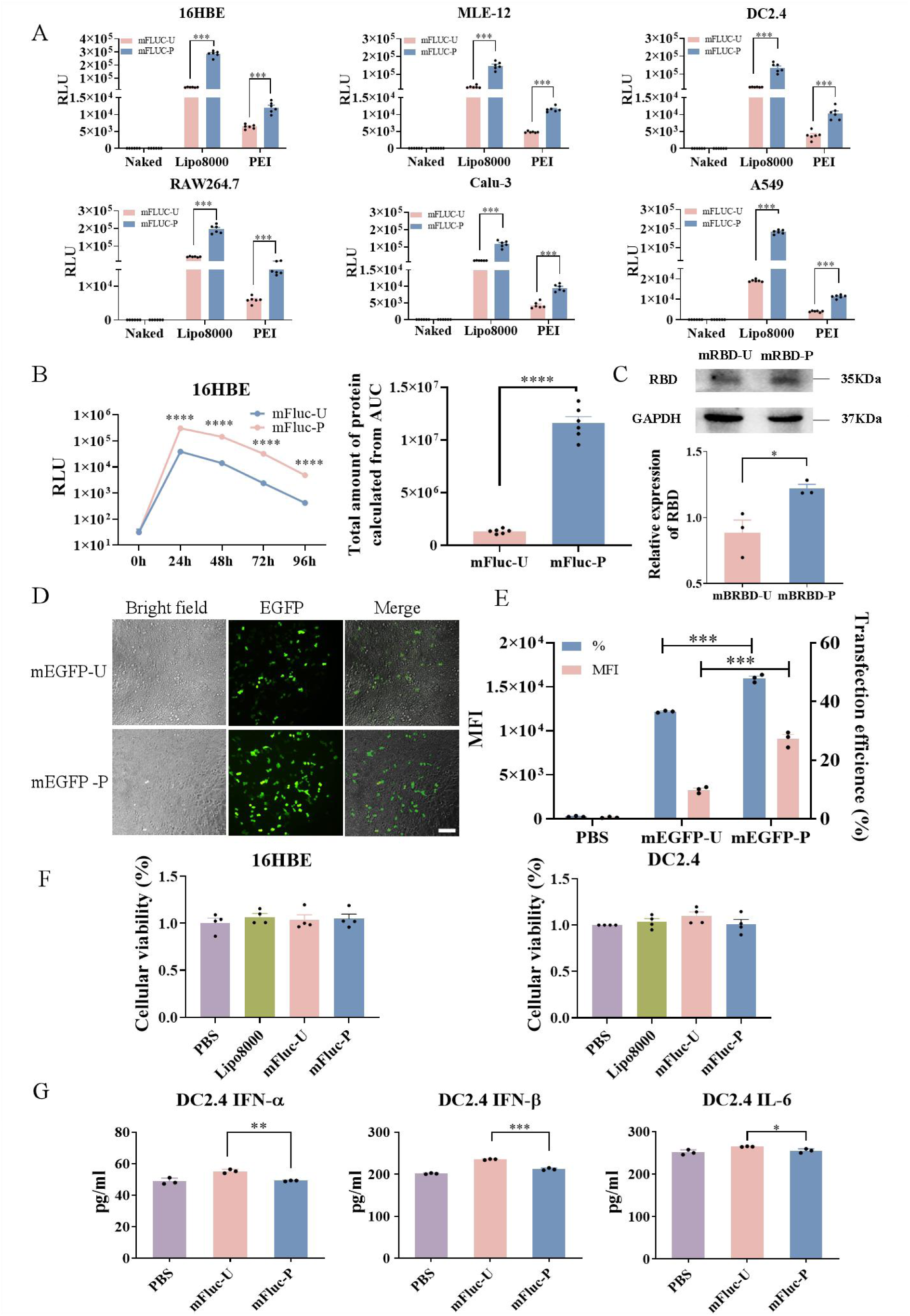
Purified IVT mRNA displays improved protein expression and safety profiles in cultured cells. (**A**) Transfection efficiency of mFluc before (mFluc-U) and after (mFluc-P) purification across various types of cultured cells, including 16HBE, MLE-12 (mouse lung epithelial cells), DC2.4, RAW264.7, Calu-3 (human lung adenocarcinoma cells) and A549. “Naked” represents naked mFluc-P, “Lipo8000” means Lipo8000-based formulations, and “PEI” represents polyethyleneimine-based formulations. (**B**) Fluc expression profile from unpurified mFluc (mFluc-U) or purified mFluc (mFluc-P) within 16HBE cells 24 h, 48 h, 72 h, 96 h post-transfection (left panel), with a corresponding quantitative comparison of the total amount of expressed protein calculated from the area under the curve (right panel). (**C**) Western blot analysis of RBD protein expression 24 h post-transfection of mRBD before (mRBD-U) and after purification (mRBD-P) (upper panel), with a semi-quantitative evaluation (n = 3 independent experiments) of the relative amount of RBD protein in western blot analysis (lower panel). (**D**) Representative transfection profiles of unpurified mEGFP (mEGFP-U) or purified mEGFP (mEGFP-P) in 16HBE cells 24 h post-transfection. Images were obtained by fluorescence microscopy with equal acquisition parameters. Scale bar, 100 μm. (**E**) Flow cytometry analysis of mEGFP-U and mEGFP-P induced transfection efficiency in 16HBE cells. (**F**) CCK8 assay evaluating the impact of purification on the viability of 16HBE and DC2.4 cells 24 h post-transfection with empty Lipo8000 without mRNA (Lipo8,000), mFluc-U and mFluc-P. (**G**) ELISA quantification of IFN-α, IFN-β and IL-6 secreted by DC2.4 cells 24 h post-transfection of mFluc-U and mFluc-P. Statistical significance was assessed by two-sided unpaired *t*-test (\**P* < 0.05; \*\**P* < 0.01; \*\*\**P* < 0.001; \*\*\*\**P* < 0.0001). The results of C and D are representative images from at least three independent experiments with similar results.

### 5. Purified IVT mRNA boosts protein expression in vivo, especially in the airway

To investigate the in vivo performance of purified-versus unpurified-IVT mRNA, mFluc-P or mFluc-U were encapsulated into LNPs and administered to mice via i.m. or i.h. routes. As shown in Fig. 5A, both purified and unpurified mRNA exhibited robust and sustained protein expression over 120 h when applied via the i.m. approach, with purified mFluc (i.m.-P) showing a 3-fold increase than the unpurified (i.m.-U) counterpart (Fig. 5B). On the other hand, the necessity and benefits of purification become distinctly more critical in terms of i.h. administration. The purified mFluc (i.h.-P) demonstrated an almost 30-fold increase in Fluc expression compared to unpurified mFluc (i.h.-U) when administered via i.h. route, with prolonged duration of protein expression up to 96 h post-dosing (Fig. 5B). This stark contrast between i.h. and i.m. routes highlights that respiratory administered IVT mRNA critically depends on its purity to achieve optimal protein expression while the intramuscularly delivered counterpart is more tolerant to the mRNA contaminants for efficient transfection. To further clarify the tissue distribution of successfully transfected mFluc, we characterized the Fluc expression in various organs of mice 6 h after dosing with mFluc-U or mFluc-P via i.m. and i.h. routes. Bioluminescence signals of mice treated via the i.m. route primarily localized in the liver and spleen, whereas only the lung displayed observable signals in mice administered via i.h. route (Supplementary Fig. 2). Apart from an improvement in transfecting the livers of mice, the i.m.-P failed to show significantly enhanced mRNA transfection compared to i.m.-U in terms of live mice and spleens (Fig. 5C). The Fluc expression detected from live mice as well as excised lungs of i.h.-P group was considerably higher than i.h.-U counterpart (Fig. 5D). All these data indicate that high-quality IVT mRNA with removal of contaminants is essential for respiratory tract-applications while the protein expression of i.m. injected mRNA seems been less influenced by the purification process.

**Fig. 5.**
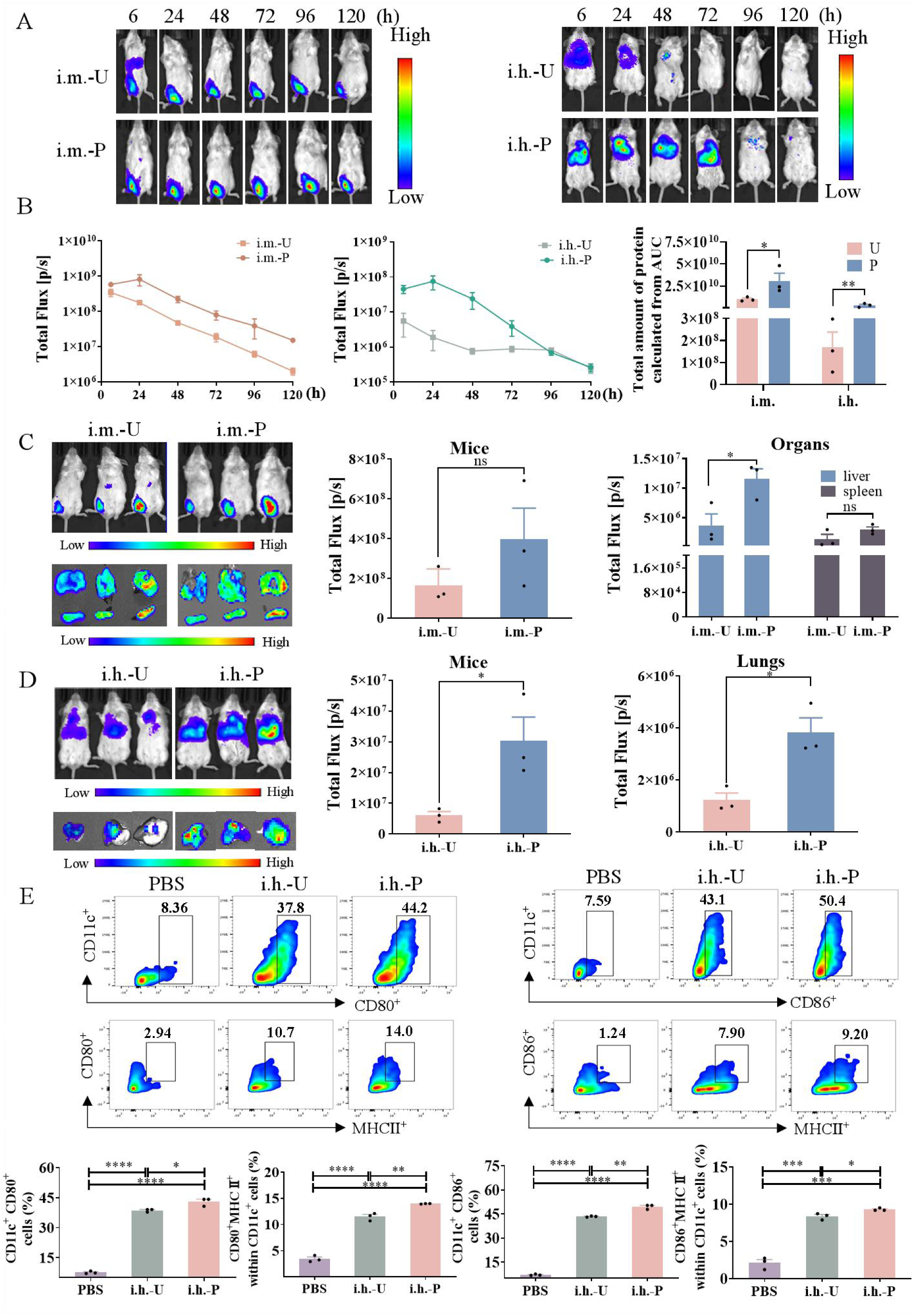
In vivo evaluations of the optimized IVT mRNA before and after purification. (**A**) Bioluminescence imaging of BALB/c mice treated with mFluc-U or mFluc-P at indicated time points post intramuscular (i.m.-U and i.m.-P) or inhalation (i.h.-U and i.h.-P) routes dosing. (**B**) Quantitative analysis of the bioluminescence signal from i.m.-U/i.m.-P and i.h.-U/i.h.-P over time and the total amount of Fluc expression was calculated from the area under curve (AUC) and compared. n = 3 biologically independent samples. (**C, D**) Representative bioluminescence images of mice and excised organs (left panel) 6 h post transfection of i.m.-U/i.m.-P (C) or i.h.-U/i.h.-P (D), with corresponding quantitative comparisons of the bioluminescence signals from live mice or excised organs (middle and right panel). (**E**) The expression of CD11c^+^CD80^+^ and CD80^+^MHC-II^+^ (gated on CD11c^+^) or CD11c^+^CD86^+^and CD86^+^MHC-II^+^ (gated on CD11c^+^) on BMDC 24 h post administrated with unpurified mRBD (i.h.-U) or purified mRBD (i.h.-P) based formulations, phosphate buffered saline (PBS) treated samples were severed as negative controls. Statistical significance was assessed by two-tailed unpaired *t*-test and one-way ANOVA with Dunnett’s post-hoc test (ns: not significant; \**P* < 0.05; \*\**P* < 0.01; \*\*\**P* < 0.001; \*\*\*\**P* < 0.0001).

Considering mature dendritic cells (DCs) are powerful antigen-presenting cells (APCs) and play key roles in initiating antigen-specific immune responses for mRNA vaccines(*39*), we additionally evaluated the ability of mRBD-P and mRBD-U based formulation in promoting DCs maturation. Compared to unpurified counterpart (i.h.-U), purified mRBD (i.h.-P) could induce higher levels of CD11c^+^CD80^+^ expressions and CD11c^+^CD86^+^ expressions on bone marrow derived dendritic cells (BMDCs) (Fig. 5E). Apart from that, the expressions of CD80^+^MHCII^+^ and CD86^+^MHCII^+^ on CD11c^+^ BMDCs were significantly elevated as well (Fig. 5E), implying the purified mRNA are more effectively in promoting the maturation of BMDCs.

### 6. Purified IVT mRNA contributes to reduce in vivo side-effects caused by dsRNA in the airway

We next investigated the impacts of purified IVT mRNA on the safety profiles of treated mice. 24 h post dosing with unpurified mRBD (U), purified mRBD (P), PBS and an empty LNP formulation without IVT mRNA (Empty) via i.m. or i.h. routes, no sign of significant necrosis, edema, inflammation or neutrophilic infiltration was detected in lung sections of the i.h.-P group, whereas mild peribronchial inflammation could be occasionally observed in lung sections of the i.h.-U group (Fig 6A). Notably, this trend was not shown in the i.m. groups, nor were differences in inflammation noted in other major organs, like livers and spleens, in i.m. or i.h. group, further observation of the kidneys, hearts, and brains of i.h.-P group showed no inflammation compared to the PBS group (Supplementary Fig. 4A). The complete blood chemistry metrics further demonstrated that the purification process could mitigate the elevation of alanine aminotransferase (ALT) and aspartate aminotransferase (AST) which are indicators of liver toxicity, this trend was observed in both i.m. and i.h. groups, although no significant differences were discernible between U and P (Supplementary Fig. 4B). To further investigate the activation of the innate immune response(*40*), the lung tissues from the i.h. group were analyzed for the expression levels of some representative cytokines, including IFN-α, IFN-β, and interleukin-6 (IL-6). All these cytokines display a significant rise in the i.h.-U group compared to both PBS and Empty controls, while the cytokine levels in the i.h.-P group were similar to the controls (Fig. 6B), suggesting a low innate immune stimulating profile of purified IVT mRNA.

**Fig. 6.**
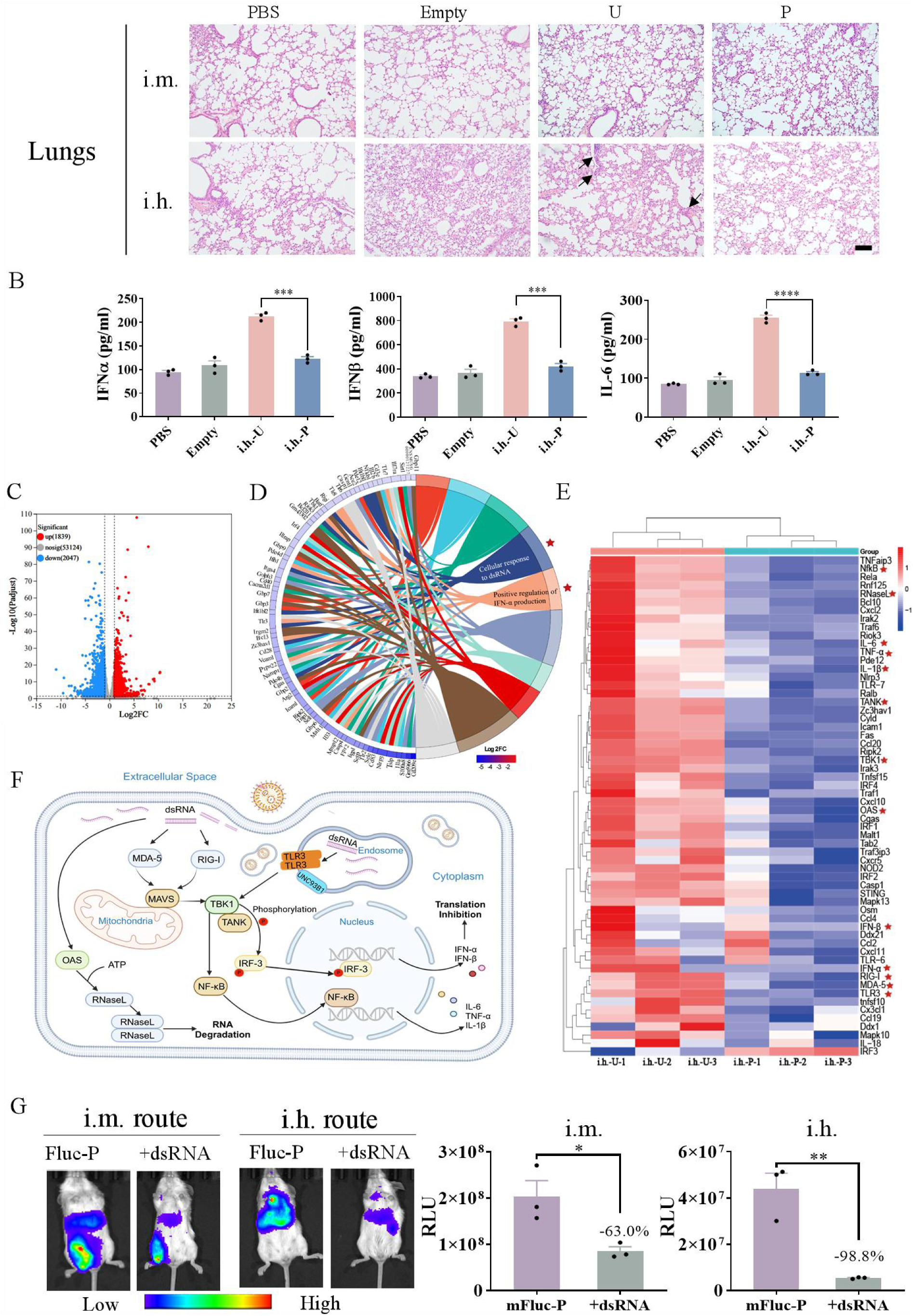
Evaluations of IVT mRNA purification on in vivo toxicity and devastating impacts of dsRNA contaminant. (**A**) Representative hematoxylin and eosin (H&E) staining of lung sections collected 24 h after administration (i.m. and i.h.) of unpurified mRBD (U) and purified mRBD (P). Samples treated by PBS and LNP formulation without mRNA (Empty) via i.h. route were used as controls. Scale bar, 100 μm. (**B**) Analysis of cytokines secretion in mice dosed by PBS, Empty, mRBD-U and mRBD-P via i.h. route. Lungs were collected 24 h post administration and IFN-α, IFN-β and IL-6 were measured by ELISA. (**C**) Volcano plot representing the upregulated and downregulated genes. (**D**) Chord plot depicting the relationship between downregulated inflammation-related genes and KEGG pathways. (**E**) Heat map representing the downregulated cytokines, chemokines and receptors. (**F**) Schematic illustration on pathways of dsRNA-induced inflammatory responses. Created using BioRender.com. (**G**) Representative bioluminescence images of mice (left panel) treated by purified mFLuc (mFluc-P) or purified mFLuc with the addition of 1.0% dsRNA (+dsRNA, w/w) via i.m. or i.h. routes (left panel) 6 h post-transfection, with corresponding quantitative comparisons of the bioluminescence signals (right panel). Each symbol represents one animal (n = 3 biologically independent mice). Statistical significance was assessed by two-tailed unpaired *t*-test (\**P* < 0.05; ** *P* < 0.01; *** *P* < 0.001; *** *P* < 0.0001).

The superiority of purified mRBD was further revealed by an RNA sequencing analysis of lung tissues from the i.h. group. A total of 3,886 differentially expressed genes were identified, including 1,839 up-regulated and 2,047 down-regulated genes (Fig. 6C). KEGG enrichment analysis showed that the downregulated genes related to inflammation were significantly enriched in dsRNA signaling pathway and the pathways associated with the production of IFN-α (Fig 6D), underscoring the purified mRNA tends to avoid excessively activating the innate immune responses due to its weak dsRNA stimulation. Key mediator genes which represent crucial sensors of dsRNA within these pathways, including *TLR-3* (T*oll-Like Receptor 3*), *RIG-I* (*Retinoic Acid-Inducible Gene I*), *MDA-5* (*Melanoma Differentiation-Associated protein 5*), and *TBK1* (*TANK-binding kinase 1 Gene*) were significantly down-regulated in lung tissues of i.h.-P (Fig 6E). Upon dsRNA recognition, TLR-3, RIG-I and MDA-5 trigger a cascade of signaling events through activating TBK1 (TANK-binding kinase 1) (*41*). The activated TBK1 leads to the activation of downstream transcription factors NF-κB (Nuclear Factor Kappa B) and IRF3 (Interferon Regulatory Factor 3) for producing type I interferons and other proinflammatory cytokines, such as IFN-α, IFN-β, IL-6, IL-1β and TNF-α. In addition, dsRNA potentially activates the up-regulation of *OAS* and down-stream *RNase L* (Ribonuclease L), which stimulates RNase L production and cleaves exogenous RNA (including the administered IVT mRNA)(*33*) (Fig 6F). The purified IVT mRNA prominently reduced the activation levels of these “notorious” genes and the inflammatory pathways mentioned above in the treated mice, indicating the efficient removal of dsRNA due to purification significantly decreases the activation of dsRNA sensors, thus curtailing the release of inflammatory cytokines, type I interferons and the activation of exogenous IVT mRNA degradation mechanism represented by the downregulated *RNase L* gene(*33, 42*).

To further examine the negative impact of dsRNA on IVT mRNA, we incorporated a 1% (w/w) reference standard of dsRNA into the purified IVT mRNA and transfected mice via i.m. and i.h. routes. Results indicate that the addition of dsRNA caused a slight increase in the particle size of the LNP formulation; no differences were observed in the PDI, zeta potential, or encapsulation efficiency of the formulation (Supplementary Fig. 5A-5D). However, the dsRNA significantly impairs the mRNA expression in both dosing approaches, but this detrimental effect varies a lot between i.m. and i.h. routes (Fig. 6G). The presence of dsRNA (+dsRNA) leads to an average 63.0% reduction of mRNA transfection in the i.m. group compared to the control (mFluc-P), whereas 98.8% of bioluminescence signal of mice transfected via i.h. route has vanished once dsRNA was included (+dsRNA). This dichotomy in i.m. and i.h. approaches is, in part, due to the existence of a dense network of immune surveillance mechanisms in the airway, which results in a more sensitive innate immune activation to dsRNA, even in a very limited amount(*43*). We infer receptors such as *TLR3*, *RIG-I*, *MDA-5* and *OAS* are more actively engaged in the lung tissue, leading to a hyperactive innate immune activation that severely compromises IVT mRNA stability and translation efficiency. These findings emphasize that purification is critical to minimize dsRNA content and maximize the potency of airway-applied mRNA therapeutics.

### 7. Purified mRNA vaccine substantially improved humoral and mucosal immune responses when inoculated via the airway

To determine the impact of purification on the immunogenicity of mRNA vaccine, mice were immunized with mRBD-U or mRBD-P via i.m. or i.h. routes using an immunization scheme as shown in Fig. 7A. The mRBD-P inoculated via i.h. route (i.h.-P) substantially increased RBD-specific IgG antibody in serum samples compared to unpurified counterpart (i.h.-U) at 14 days and 28 days post prime with 3 μg mRBD/mouse (Fig. 7B). Whereas mice vaccinated by mRBD-P (i.m.-P) and mRBD-U (i.m.-U) through i.m. route did not show substantial differences during the investigated period (Fig. 7B). We assume this phenomenon is caused by the different mRNA expression profiles between i.h. and i.m. routes, because the efficacy of mRNA vaccine is closely linked to antigen expression level. The i.m. administered mRNA vaccine (irrespective of purified or unpurified ones) could relatively easier translate enough amount of antigens to activate the immune system when inoculated with a high-dose of mRNA (3 μg mRBD/mouse). As a result, we further evaluated the impact of purification when a low-dose of mRNA vaccine (1 μg mRBD/mouse) was applied. Interestingly, both low-dose i.h.-P and i.m.-P increased antigen-specific IgG antibodies in serum samples on day 14 and day 21. On day 28, the i.h.-P group showed significantly higher IgG levels compared to the i.h.-U group. However, the i.m.-U and i.m.-P groups had comparable levels of RBD-specific IgG antibodies at the same time point after the boost dose (Fig. 7C). These data confirmed our hypothesis and suggest the different antigen expression profiles between low-dose i.m.-P and i.m.-U could be compensated by the boost dose.

**Fig. 7.**
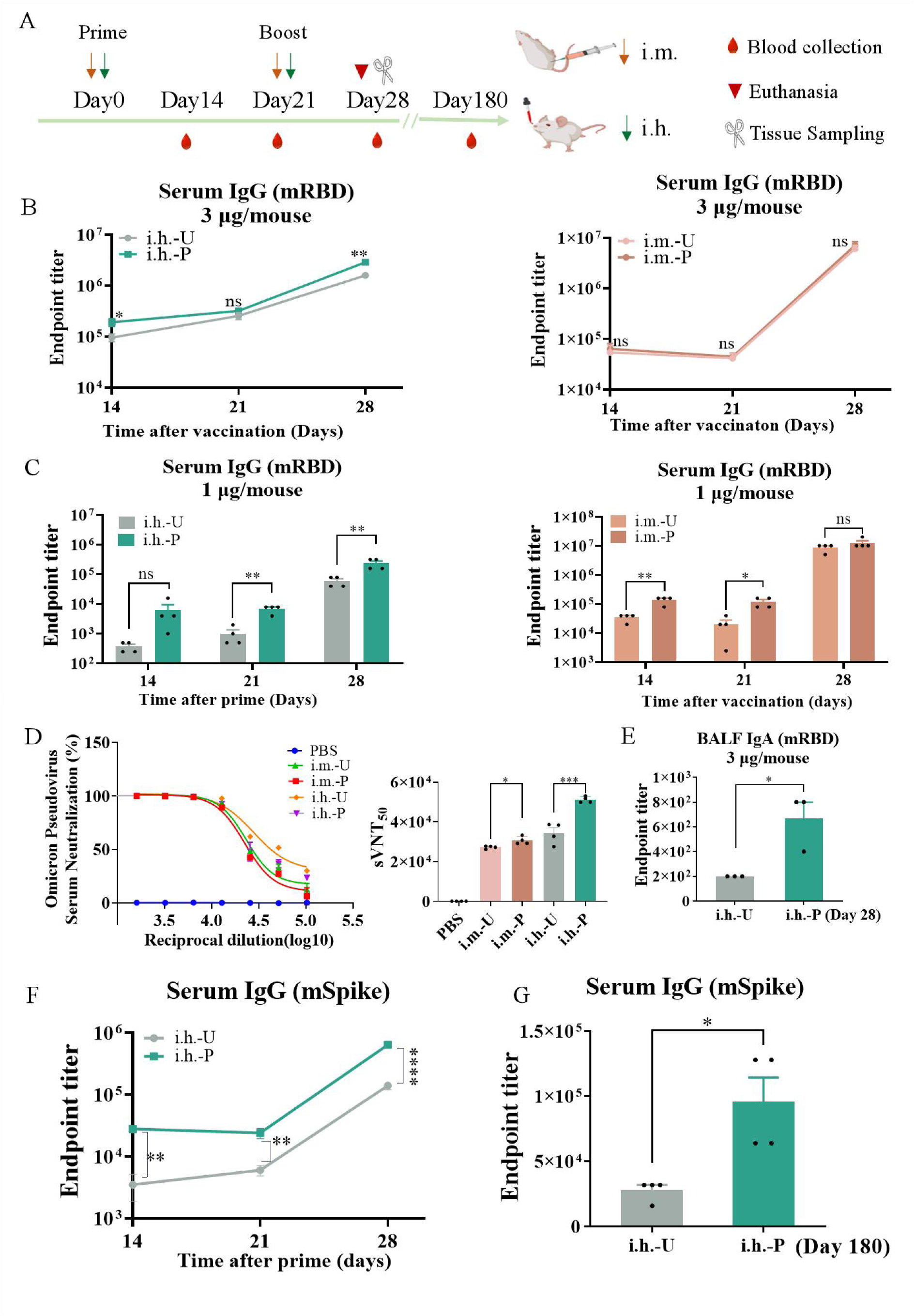
Humoral and mucosal immune responses induced by IVT mRNA with or without purification in mice inoculated via i.m. or i.h. routes. (**A**) Schematic of immunization and sampling procedure. (**B**) Endpoint titers of RBD-specific IgG antibody in serum samples collected at 14 days, 21 days and 28 days post prime with LNPs containing 3 μg/mouse mRBD-U or mRBD-P via i.m. or i.h. routes (the boost dose on day 21 was the same as the prime dose), n = 4 biologically independent samples. (**C**) Endpoint titers of RBD-specific IgG antibody in serum samples collected at indicated time points after prime and boost with LNP encapsulating 1 μg/mouse mRBD-U or mRBD-P via i.m. and i.h. routes, n = 4 biologically independent samples. (**D**) Pseudovirus neutralizing antibody titers in serum samples were collected on day 28 using the same dosing scheme as described in (B). sVNT_50_, Serum Virus Neutralization Test 50% endpoint, n = 4 biologically independent samples. (**E**) Endpoint titers of RBD-specific sIgA antibody in BALF samples were collected on day 28 using the same dosing scheme as described in (B), n = 3 biologically independent samples. (**F**) Endpoint titers of SARS-CoV-2 spike-specific IgG antibody in serum samples at 14 days, 21 days and 28 days post prime and boost with LNPs containing 3 μg/mouse mSpike-U or mSpike-P via i.h. routes, n = 4 biologically independent samples. (**G**) Endpoint titers of spike-specific IgG antibody in serum samples collected on day 180 using the same dosing scheme as described in (F), n = 4 biologically independent samples. Statistical significance was calculated by two-tailed unpaired *t*-test (ns: not significant; \**P* < 0.05; \*\**P* < 0.01; \*\*\**P* < 0.001; \*\*\*\**P* < 0.0001).

Pseudovirus neutralization assays further reveal that both i.h.-P and i.m.-P significantly enhanced neutralizing antibody titers against the Omicron variant compared to their unpurified counterparts (Fig. 7D), but with a more pronounced improvement in the i.h. group (i.e. i.h.-P VS i.h.-U) in which the serum Virus Neutralization Test 50% endpoint (sVNT_50_) of i.h.-P approached ∼1/50000. Apart from the enhanced serum IgG antibody, the i.h.-P can significantly enhance the mucosal immune responses as well. The RBD-specific sIgA antibody in BALF samples collected from i.h.-P reached an endpoint titer up to 1/800 and was significantly higher than i.h.-U counterparts (Fig. 7E). Besides, compatibility of mRNA with different lengths is an important criterion for affinity chromatography purification(*44*). In order to investigate the impact of purification on the immune efficacy of mRNA with longer sequences, unpurified (U) and purified (P) IVT mRNA encoding SARS-CoV-2 spike protein (mSpike), which has a length of approximately 4,000 nt, was selected and vaccinated mice via i.h. route. Consistently, mSpike-P displayed higher levels of SARS-CoV-2 spike-specific IgG antibody in serum samples compared to mSpike-U counterpart (Fig. 7F), and this trend persisted for at least 180 days (Fig. 7G). The above data indicate that purification has positive impacts on the immunogenicity of mRNA vaccines with both short and long sequences. Purified mRNA vaccines are more efficient than unpurified counterparts in promoting efficient and long-lasting antigen-specific humoral and mucosal immune responses when inoculated via the respiratory route. In contrast, the purification process poses fewer benefits to the mRNA vaccines administrated via i.m., which presumably resulted from a similar antigen expression profile between the purified and unpurified mRNA.

### 8. Purification substantially enhances cellular immune responses induced by mucosal mRNA vaccines

To further explore the cellular immune responses induced by mRBD-U and mRBD-P via i.m. or i.h. administration, we measured IFN-γ, IL-4, and IL-17A production in pulmonary lymphocytes and splenocytes of vaccinated mice using the ELISpot assay. As shown in Fig. 8A, pulmonary lymphocytes from the i.h.-P and i.m.-P groups contained significantly more IFN-γ and IL-4 secreting cells than their respective unpurified counterparts (i.e. i.h.-U and i.m.-U). Additionally, a significant increase in IL-17A secreting cells was only observed in the i.h.-P group, with no notable differences in the pulmonary lymphocytes between the i.m.-U and i.m.-P groups (Fig. 8A). A similar pattern was seen in splenocyte samples, indicating that mRNA purification boosts antigen-specific cytokine production in both lungs and spleens (Fig. 8B). We then assessed intracellular IFN-γ production in CD4^+^ and CD8^+^ T cells using flow cytometry, it reveals a notable improvement in IFN-γ secretion within pulmonary lymphocytes from both i.m.-P (Fig. 8C) and i.h.-P groups (Fig. 8D). However, this trend was only observed in CD8^+^IFN-γ^+^ splenocytes from the purified groups (i.e. i.m.-P and i.h.-P), with no significant change detected in CD4^+^IFN-γ^+^ splenocytes (Fig. 8E and 8F). It is worth noting that CD8^+^IFN-γ^+^ cells within pulmonary lymphocytes and splenocytes collected from the i.h.-P group (Fig. 8D and 8F) demonstrated a qualitative change from background level to significantly detectable amounts, underscoring the profound impact of purified mRNA in eliciting a comprehensive and potent cellular immune response, particularly when administered via i.h. route.

**Fig. 8.**
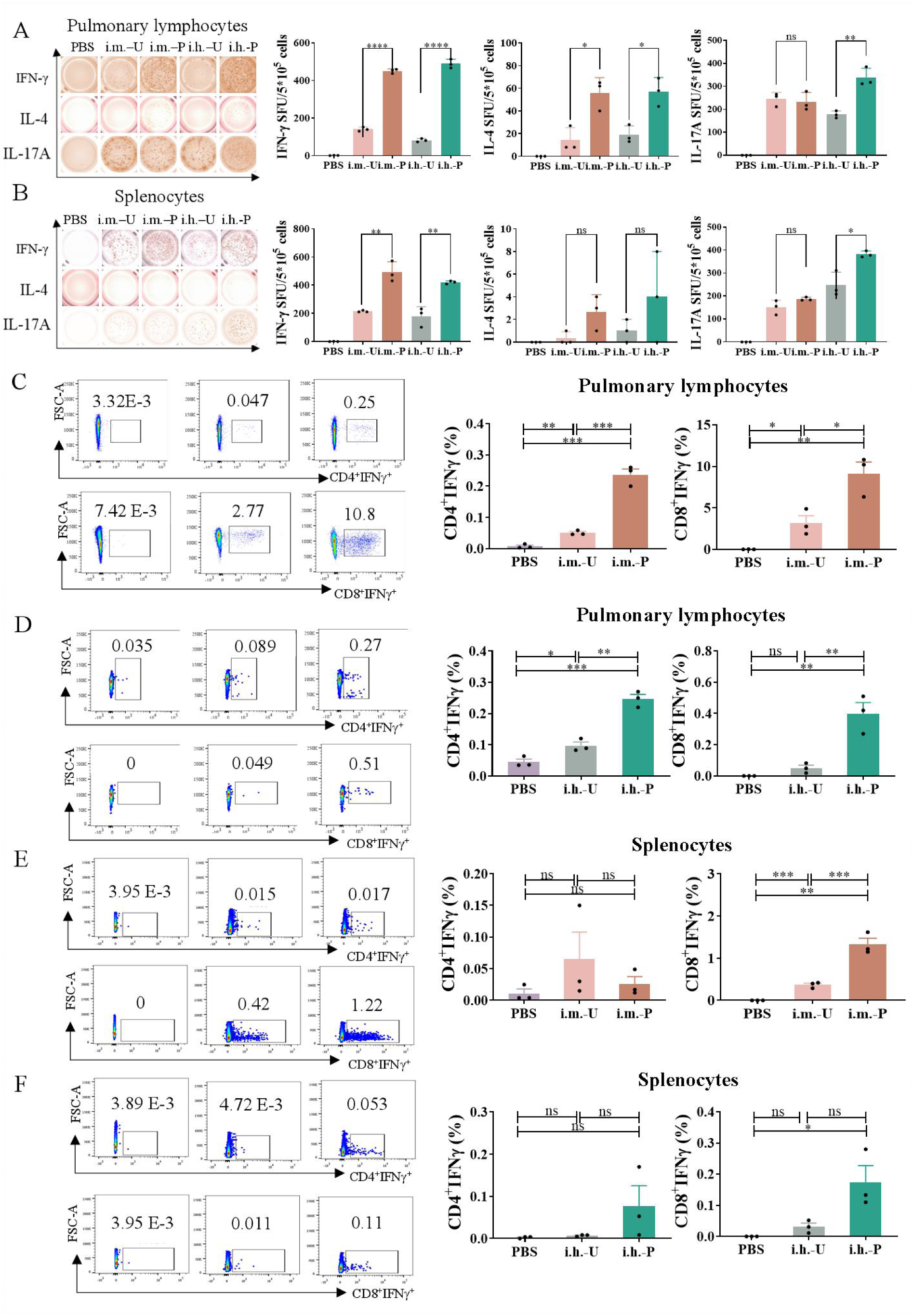
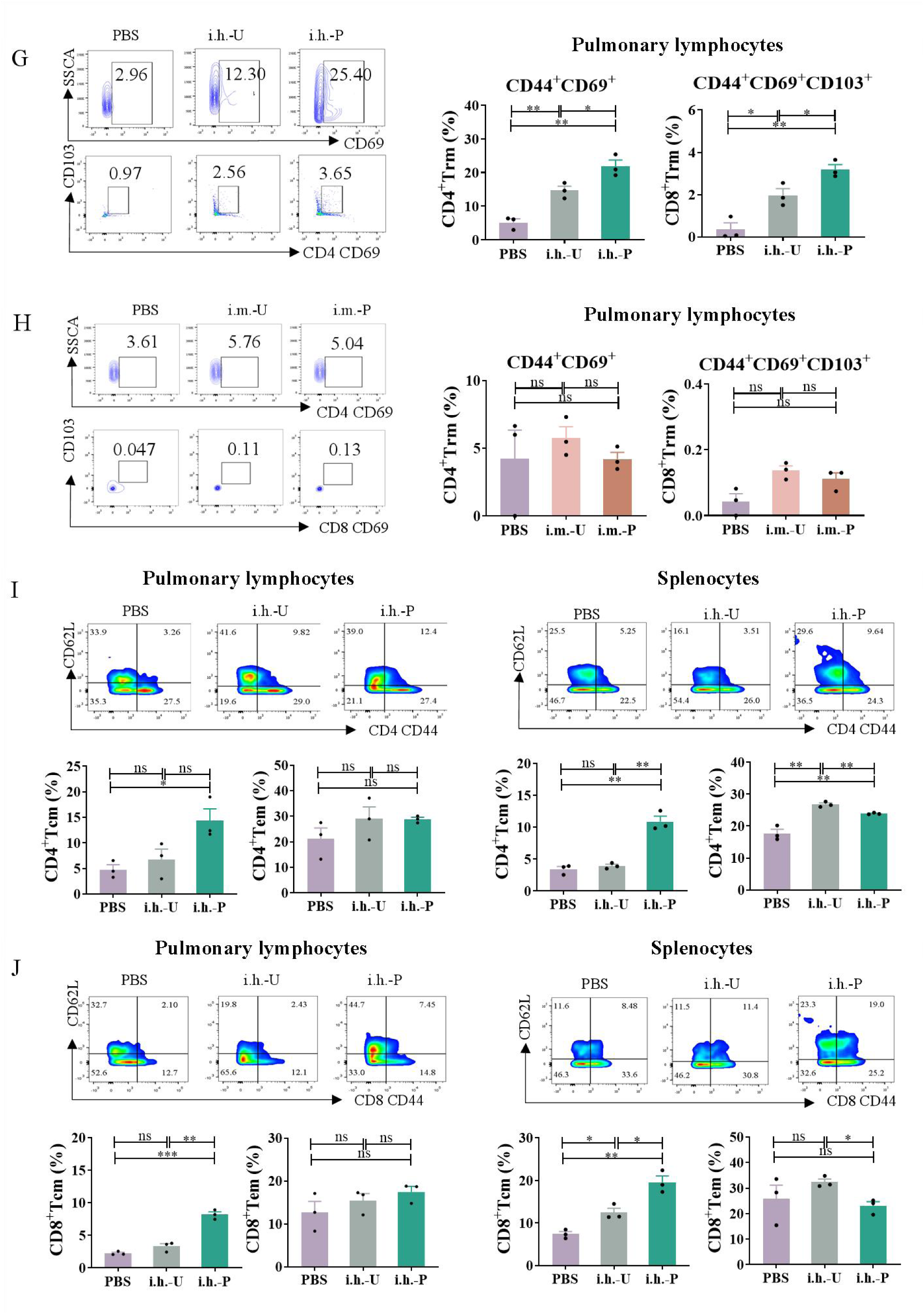
Immunologic evaluations of SARS-CoV-2 specific T cell responses and memory-biased immunity post-immunization with unpurified mRBD (U) or purified mRBD (P) via i.m. or i.h. routes. (**A, B**) IFN-γ, IL-4, and IL-17A producing cells being quantified by the ELISpot assay using pulmonary lymphocytes (A) and splenocytes (B) re-stimulated with SARS-CoV-2 RBD peptide pool (details can be found in Supplementary Table 6). (**C, D**) Flow cytometry analysis on the proportion of RBD-specific intracellular cytokine of IFN-γ secreting CD4^+^ and CD8^+^ T cells within RBD peptide pool re-stimulated pulmonary lymphocytes collected from mice vaccinated via i.m. (C) or i.h. (D) routes. (**E, F**) Measurement of IFN-γ secreting CD4^+^ and CD8^+^ T cells within splenocytes generated from i.m. counterparts (E) or i.h. counterparts (F) treated in the same way as described in C, D. (**G, H**) Trm cells frequencies across CD4^+^ (CD4^+^CD44^+^CD69^+^) and CD8^+^ (CD8^+^CD44^+^CD69^+^CD103^+^) T cells in the lungs of mice vaccinated via i.m. (G) or i.h. (H) routes. (**I, J**) Levels of Tcm (CD44^+^CD62L^+^) and Tem (CD44^+^CD62L^-^) across CD4^+^ and CD8^+^T cells in the lungs (I) or spleens (J) of mice vaccinated via i.h. routes. Each symbol in A-J represents one biologically independent animal. Statistical significance was calculated by one-way ANOVA with Dunnett’s post-hoc test (ns: not significant; \**P* < 0.05; \*\**P* < 0.01; \*\*\**P* < 0.001; \*\*\*\**P* < 0.0001).

Compelling evidences indicate the crucial roles of lung Trm cells in eliminating SARS-CoV-2 and its viatants of concern(*45, 46*). To determine how the purified mRNA vaccine impact on the memory T cell responses within pulmonary mucosal sites, we investigated the CD4^+^ Trm cells (CD4^+^CD44^+^CD69^+^) and CD8^+^ Trm cells (CD8^+^CD44^+^CD69^+^CD103^+^) cells isolated from the lungs of mice immunized via i.m. or i.h. routes. Results suggested that both the CD4^+^ Trm and CD8^+^ Trm cells in the i.h.-U group were significantly enhanced compared to PBS-treated counterparts and this improvement could be further substantially enlarged when mice were vaccinated by i.h.-P (Fig. 8G). However, the amount of pulmonary CD4^+^ Trm and CD8^+^ Trm cells in the i.m.-U. and i.m.-P groups were in a comparable level which is similar to PBS-treated counterparts (Fig. 8H). We also evaluated the CD4^+^ Trm cells and CD8^+^ Trm cells in the spleens of mice immunized via i.m. or i.h. routes but unfortunately obtained negative results in all tested groups (Supplementary Fig. 6A), indicating the Trm cells are not established in the spleen. Effector memory T (Tem) cells and central memory T (Tcm) cells, identified as CD44^+^CD62L^-^ and CD44^+^CD62L^+^, were also detected from the pulmonary or splenic samples of vaccinated mice. Intriguingly, we found i.h.-P vaccination consistantly boosted the ratio of Tcm (both CD4^+^ and CD8^+^ based) cells compared to i.h.-U and PBS counterparts, no matter in the samples collected from the lung or the spleen, whereas i.h.-P vaccination did not show any advantages in terms of enhancing CD4^+^ Tem and CD8^+^ Tem cells (Fig. 8I and 8J). Similar to what we observed in Trm cells, purified mRNA (i.m.-P) did not have benefits either on the Tem cells or the Tcm cells when inoculated via the i.m. route (Supplementary Fig. 6B and Supplementary Fig. 6C). These observations denote the tremendous potential of purified mRNA in inducing comprehensive mucosal T cell immune responses and memory-biased immunity in the vaccinated candidates.

## Discussion

In this study, we demonstrate that the intrinsic quality of IVT mRNA, particularly the sequence design and purity, is crucial for its effectiveness when administered via the respiratory route (Supplementary Fig. 7A). Notably, the airway-administered mRNA vaccines require higher standards of purification and tailor-designed UTRs. This is reflected in the enhanced mRNA transfection efficiency in which the optimized mRNA could mediate up to 30-fold total protein expression when dosed via the i.h. route, in contrast to the less than 4-fold improvement when the same mRNA was inoculated via the i.m. route (Supplementary Fig. 7B-7D). The enhanced target antigen expression subsequently enables the airway-administered mRNA outperform parenteral injected counterparts in mediating more potent adaptive immune responses, including humoral immunity (Supplementary Fig. 7E-7G), cellular immunity (Supplementary Fig. 7H-7M), and mucosal immunity (Fig. 8E). This phenomenon most likely arises from the insurmountable mucociliary clearance mechanism and sophisticated immunological environment of respiratory mucosa, which have evolved over millions of years to adeptly remove foreign substances(*43, 47*). Intriguingly, while minor impurities in parenteral route inoculated mRNA typically do not markedly affect vaccine efficacy, they can precipitate innate immune reactions within the respiratory system. We identified such side effects are primarily mediated through the recognition of dsRNA as a pathogen-associated molecular pattern. This recognition is facilitated by innate immune receptors such as TLR3, RIG-I, MDA-5, and the activation of RNase L(*30*). Engagement of these receptors leads to the activation of complex signaling pathways, including the RIG-I, MDA-5 and OAS pathways, culminating in the rapid production of type I interferons and other pro-inflammatory cytokines. This cascade results in the swift clearance of foreign IVT mRNA in the airway, substantially reducing the expression of the target antigen and thereby attenuating the potential for a robust adaptive immune response.

To address these challenges, we developed a simple and mild IVT mRNA purification approach based on affinity chromatography to remove the unwanted culprits within IVT mRNA. This purification process effectively removes contaminants such as dsRNA. The primary advantages of using affinity chromatography include its ability to operate under mild conditions, which preserves the integrity and functionality of IVT mRNA with high yields (achieving up to 80% recovery rate under optimized chromatographic conditions) and purity suitable in cGMP production, and its scalability, which is vital for mass production of mRNA vaccines(*48*). By utilizing highly pure IVT mRNA, these vaccines can achieve higher stability and improved effectiveness in the respiratory tract, where the presence of impurities could otherwise lead to rapid degradation of IVT mRNA and undetectable levels of antigen-specific immune responses. Improving the purity of IVT mRNA through affinity chromatography is essential for developing mucosal vaccines that may provide robust protection against airborne pathogens.

Our strategies in ameliorating pulmonary applied IVT mRNA further include a rational design of UTRs sequence, which is specifically tailored for respiratory applications. The UTRs have been screened out from a large set of genes that are highly expressed in pulmonary cells. Based on the UTRs originating from nature, specialized elements (such as the Apt-eIF4G and R3U) have been incorporated to further enhance transcriptional output and translation initiation. Modifications at the 3’ UTR, such as the removal of miRNA binding sites, are crucial to prevent post-transcriptional downregulation, thereby extending the stability and effectiveness of IVT mRNA within the respiratory tract. Not only enhance the stability and translation efficiency of IVT mRNA by these modifications in the target cells, but also the optimized UTRs contribute to evading rapid degradation(*49*). This specialized sequence optimization ensures that the mRNA remains functional for a longer period within the aggressive environment of the respiratory tract, leading to a more sustained and potent immune response.

In summary, this study demonstrates that the quality of IVT mRNA itself is critically important for its efficacy when applied via the respiratory route. The dsRNA contaminants within IVT mRNA are prone to lead to a hyperactive innate immune activation in the airway that severely compromises the transfection efficiency of IVT mRNA. Our findings emphasize the critical need for high-quality mRNA molecules for mucosal mRNA applications, which should ideally incorporate sequences specifically designed for the airway route and exhibit high purity with minimal contaminants. Top-notch mRNA molecules tend to expedite the clinical translation process of the next-generation mucosal mRNA vaccines that hold the ability to control invading airborne pathogens at the initial port of infection and potentially enable us to be better prepared in combating the outbreak of pandemic in the future.

## MATERIALS AND METHODS

### Reagents

Dulbecco’s Modified Eagle Medium (DMEM), streptomycin and Trypsin-EDTA were purchased from Gibco. Opti-MEM I Reduced Serum Medium, calcium chloride (CaCl_2_), magnesium chloride (MgCl_2_), nuclease-free water and Lipofectamine3,000 were purchased from Invitrogen Thermo Fisher Scientific. Phosphate buffered saline (PBS) was purchased from Biosharp. Na₃PO₄, NaCl and NaOH were purchased from Sigma-Aldrich. Agarose was purchased from Aladdin, diethylpyrocarbonate (DEPC) was purchased from Beyotime. 3-Morpholinopropanesulfoinc Acid (MOPS) buffer and formaldehyde were purchased from Solarbio. Lipo8,000 was purchased from Beyotime and polyethyleneimine (PEI) was purchased from MKBio. RIPA buffer was purchased from Cell Signaling Technology. A 1% protease inhibitor cocktail was purchased from Roche. PVDF membrane was purchased from Millipore. Tween 20 was purchased from AppliChem. Skimmed milk powder and tris buffered saline with Tween-20 (TBST) were purchased from Sbjbio and Beyotime perspectively. 4-(dimethylamino)-butanoic acid, (10Z,13Z)-1-(9Z,12Z)-9,12-octadecadien-1-yl-10,13-nonadecadien-1-ylester (D-Lin-MC3-DMA) was purchased from Shanghai Advanced Vehicle Technology L.T.D. Co (AVT). 1,2-distearoyl-sn-glycero-3-phosphocholine (DSPC), cholesterol and 1,2-dimyristoyl-rac-glycero-3-methoxypolyethylene glycol-2000 (DMG-PEG-2,000) were purchased from Avanti. Pre-rinsed dialysis bags were purchased from Viskase. All other solvents and small molecular reagents were obtained in high quality (analytical or HPLC grade) from Sigma-Aldrich.

### Cells

human bronchial epithelial cells (16HBE), mouse lung epithelial cells (MLE-12), mouse bone marrow-derived dendritic cells (DC2.4), and Human Embryonic Kidney 293 cells (HEK293T) stably expressing Human Angiotensin-Converting Enzyme 2 (hACE2) (HEK-293T-hACE2) were cultured in RPMI-1640 (Gibco), while mouse mononuclear macrophage leukemia cell (RAW264.7), human lung adenocarcinoma cell (Calu-3), and human non-small cell lung cancer cell (A549) were maintained in DMEM. Both media were supplemented with 100 U/mL penicillin, 100 μg/mL streptomycin, and 10% fetal bovine serum (FBS, Gibco) in an environment of 5% CO_2_ at 37 °C. Cells were trypsinized every 2-3 days using 0.25% Trypsin-EDTA for passaging. All cell lines were obtained from original providers who authenticated the cell lines using morphology, karyotyping and PCR-based approaches. No additional authentication has been performed. All experiments were performed on cells in the logarithmic growth phase. Bone marrow-derived dendritic cells (BMDCs) were generated from specific pathogen free (SPF) female BALB/c mice (6–8 weeks old) using standard protocols. All cell lines tested negative for mycoplasma contamination.

### In vitro transcribed messenger RNA (IVT mRNA)

IVT mRNAs encoding firefly luciferase (mFluc), enhanced green fluorescent protein (mEGFP), the receptor binding domain of ancestral SARS-CoV-2 (mRBD), and the wild-type spike protein of SARS-CoV-2 (mSpike) were synthesized from linearized plasmids using T7 RNA polymerase (T7 High Yield RNA Transcription kit, Novoprotein Scientific). The transcription process incorporated 5’ and 3’ UTRs derived from genes highly expressed in airway-related cells and included a 110-nucleotide-long poly (A) tail, interrupted by a short linker sequence (A30LA70), transcribed from the respective DNA templates. For the creation of nucleoside-modified mRNAs, uridine 5’ -triphosphate (UTP) was substituted with N1-Methyl-Pseudo-UTP (100 mM, Glycogene) during the transcription reaction. Capping of the IVT mRNAs was conducted co-transcriptionally (one-pot capping) using the CleanCap trinucleotide cap1 analogue (T7 High Yield RNA Synthesis Kit for Co-transcription, Yeasen Biotechnology) or enzymatically (enzymatic capping) with the Cap 1 Capping System (M082-01B, Novoprotein Scientific) post transcription. The IVT mRNA was heated to 65°C for 5 min to denature before the capping process. Following transcription, the mRNA was treated with DNase I and subsequently purified through lithium chloride precipitation using lithium chloride (AM9480, Sigma-Aldrich). The resulting pellet was dissolved in nuclease-free water. The mRNA concentration was measured using a Microvolume UV-Vis spectrophotometer (NanoDrop one, Thermo Fisher Scientific), with stock concentrations ranging between 1-3 μg/μL. The integrity and purity of the purified mRNA products were confirmed by gel electrophoresis.

### Affinity chromatography purification of IVT mRNA

IVT mRNA was further purified by High Performance Liquid Chromatography (HPLC) (Agilent, 1260) using a column CIMacTMOligo dT18 0.1 mL Analytical Column (C12 Linker, 2 μm channels, Sartorius). Buffer A (Loading buffer): 50 mM Na₃PO₄, 1M NaCl, pH = 7.0; Buffer B (Washing buffer): 50 mM Na₃PO₄, pH 7.0; Buffer C (Elution buffer): nuclease-free water; Buffer D (Cleaning buffer): 0.5 M NaOH. After optimizing, the purification method could efficiently purify different IVT mRNA. The column was initially equilibrated with 100% Buffer A. Subsequently, IVT mRNA was loaded and the column was run for 2 min in Buffer A. At the end of the second min, a gradient was initiated, decreasing Buffer A from 100% to 0%, while increasing Buffer B from 0% to 100%, maintaining this state from the 2.01th min to the sixth min to facilitate washing. At the 6.01th min, Buffer B was reduced to 0% while Buffer C was increased to 100%, maintained until the eighth min to elute the bound mRNA (collected into a fraction collector). At the 8.01th min, Buffer C was reduced to 0% and Buffer D increased to 100%, continuing until the tenth min to clean the system. The procedure concluded by returning to Buffer A to re-equilibrate the system. The sample loading volume was 50 μL, and the mRNA concentration was maintained at around 1μg/μL, with flowrate at 1 mL/min during the entire purification process. The eluted mRNA was collected using a fraction collector (Agilent Technology) and was subsequently concentrated using a vacuum concentrator (CV200, JM Technology) at room temperature (RT) at 1500 revolutions per minute (rpm) for 4 h to obtain the appropriate concentration of mRNA products.

### dsRNA dot blot

The elimination of dsRNA from the final IVT mRNA products was assessed using the dsRNA DIBA Kit (RD009, Novoprotein Scientific). Briefly, 200 ng or 400 ng of purified mRNA were applied to the nylon blotting membrane. The membrane was then blocked for 1 h in TBS-T buffer containing 5% (w/v) skimmed milk powder. The mouse monoclonal antibody (anti dsRNA) was diluted at 1:5,000 in 1% (w/v) milk/TBS-T buffer, and the membrane was incubated overnight under agitation at 4 ℃. The secondary antibody was diluted at 1:5,000 in 1% (w/v) milk/TBS-T buffer. The positive control contained was the dsRNA standard supplied by the kit. Dot blot was visualized on ChemiDoc TM Touch Imaging System (BIO-RAD). Blot densities were quantified using ImageJ software (National Institutes of Health).

### Gel electrophoresis

To analyze the integrity and size distribution of IVT mRNA, denaturing gel electrophoresis was performed. Briefly, a 1% agarose gel was prepared by dissolving 0.2 g agarose in 14.4 mL diethylpyrocarbonate (DEPC)-treated water. After heating until dissolved, 2 mL 10×3-Morpholinopropanesulfoinc Acid (MOPS) buffer and 3.6 mL formaldehyde were added to the mixture. After thoroughly mixing, the gel solution was poured into a casting tray. A comb was inserted to create wells, and the gel was left to solidify at RT. Once set, the gel was ready for loading samples. For sample preparation, 1 μg of IVT mRNA was mixed with denaturing loading buffer in a 1: 2 (v/v) ratio. The mixture was heated to 65°C for 5 min to denature the mRNA, then cooled on ice for 5 min before loading. The samples were loaded into the wells of the prepared agarose gel, and electrophoresis was conducted at 120 volts for 20 min in 1× MOPS running buffer. The progress of electrophoresis was monitored by the migration of the bromophenol blue dye marker. After electrophoresis, the gel was imaged under UV light using an Imager system (3500, Tanon) to capture the distribution and integrity of the IVT mRNA.

### Preparation of mRNA-LNP formulations

Lipid nanoparticles (LNP) encapsulated with IVT mRNA were produced by mixing an aqueous phase containing the mRNA with an ethanol phase comprising the lipids in a microfluidic chip device (AceNANOSOME; ACMEI LIFESCIENCE). Lipids were dissolved in ethanol at molar ratios of 50:10:38.5:1.5 for intramuscular injection (i.m.) route (25 mM D-Lin-MC3-DMA; 5 mM DSPC; 19.25 mM cholesterol; 0.75 mM DMG-PEG-2,000) or 46:23:30:1 for inhalation (i.h.) route applications. The self-assembly process of mRNA-LNP nanoparticles was realized by mixing the lipid solution with an acidic aqueous IVT mRNA solution (100 mM sodium acetate, pH 4.0) at a 1:3 volume ratio, and the nitrogen to phosphorus ratio (N/P) of 8:1 between the cationic amines in the lipids and the anionic phosphate. The LNP formulations were loaded into pre-rinsed dialysis bags with a 10-14 kDa Molecular Weight Cut-Off and dialyzed overnight at 4°C in PBS to remove free mRNA and low molecular weight impurities. The hydrodynamic diameter and the surface charge (zeta potential) of mRNA-LNP complexes were characterized by dynamic light scattering with the Zetasizer Nano ZS (Malvern Instruments, UK). Quantification of mRNA encapsulation into LNPs was performed with RiboGreen (Invitrogen) according to the manufacturer’s instructions. Transmission electron microscopy (TEM, JEOL Ltd.) was used to observe the morphology of LNPs. The LNPs were freshly prepared and dropped onto Quantifoil Holey Carbon foil (Micro Tools) to obtain the images.

### In vitro transfection of IVT mRNA in cultured cells

16HBE, MLE-12, DC2.4, RAW264.7, Calu-3, or A549 cells were seeded in 96-well plates at densities ranging from 1.5×10^5^ to 2.5×10^5^ cells/well 24 h before transfection to reach 70% confluence. Transfection was performed using the Lipofectamine3000, Lipo8000 or PEI according to the manufacturer’s protocol. IVT mRNA was diluted in Opti-MEM I Reduced Serum Medium, mixed with Lipofectamine3000, Lipo8000 or PEI, and incubated at RT for 15 min to form complexes. These complexes were then diluted to 100 μL, added to the wells and incubated at 37°C in a 5% CO_2_ humidified atmosphere for 4 h before the medium was replaced with fresh growth medium. Fluc expression was assessed 24 h post-transfection via a microplate luminescence detector (GloMax Navgator, Promega) using the Firefly Luciferase Reporter Gene Assay Kit (RG005, Beyotime). The mEGFP expression was qualitatively observed via an inverted fluorescence microscope (ECLIPSE T, Nikon) and quantitatively evaluated via flow cytometry (FACS Canto II, BD Biosciences). The data was analyzed with FlowJo software (version 10.6.2), with Mean Fluorescence Intensity (MFI) reflecting the geometric mean of positive cells.

### Western blot

Western blot analysis was performed to assess the expression of the RBD protein in vitro. Cell lysates were prepared using a cold RIPA buffer supplemented with a 1% protease inhibitor cocktail. After mixing the lysate with the supernatant of the culture medium, protein concentrations were determined using the BCA assay (Thermo Fisher Scientific). Then the proteins were separated on a 7.5% SDS-polyacrylamide gel and electrotransferred to a PVDF membrane at 120 volts for 70 min using a Mini Trans-Blot device (Bio-Rad). To block nonspecific binding, the membrane was incubated with 5% skimmed milk powder in TBST containing 0.1% Tween 20 for 1 h at RT. The RBD protein was probed with an anti-SARS-CoV-2 Spike RBD monoclonal antibody (Solarbio; K200105M) at a dilution of 1:5,000 in 5% milk/TBST for 1 h at RT. This was followed by incubation with goat anti-mouse IgG1 HRP-conjugated secondary antibody (Abcam; ab97240) at a 1:10,000 dilution in 5% milk/TBST for an additional hour at RT. Detection was carried out using a chemiluminescence method (ChemiDoc; Bio-Rad). GAPDH (14C10 Rabbit mAb; Cell Signaling Technology) was used as a loading control. Band densities were quantified using ImageJ software (National Institutes of Health).

### Cell Counting Kit 8 (CCK 8) assay

To assess the cytotoxicity of different IVT mRNA-based formulations, a CCK8 assay was performed. Briefly, 16HBE and DC2.4 cells were seeded in 96-well plates at a density of 10,000 cells per well and allowed to adhere overnight. After treatment with mFluc before or after purification loaded in Lipo8000 for 24 h, 10 μL of CCK8 solution (MedChem Express) was added to each well. The plates were then incubated at 37°C for 2 h in a humidified atmosphere containing 5% CO_2_. The absorbance at 490 nm was measured using a microplate reader (Spectra MAX190, Molecular Devices) to determine the metabolic activity of the cells, which is indicative of cell viability. The background absorbance of the medium with the CCK8 reagent but without cells was subtracted from all readings. A control group treated with PBS alone was included to normalize the results and calculate the percentage of cell viability relative to the control.

### Animal studies

All animal procedures were approved by the Laboratory Animal Welfare and Ethics Committee of Third Military Medical University (AMUWEC20230184) and were performed in accordance with the institutional and national policies and guidelines for the use of laboratory animals. Specific pathogen-free (SPF) female BALB/c mice (6-8 weeks old) were obtained from Beijing Vital River Laboratories (Beijing, China). The mice were housed under controlled conditions of temperature and humidity with a 12 h light/dark cycle in SPF facilities. The mice were provided with free access to sterile food and water. The mice were randomly assigned to experimental groups and allowed an adaptation period of at least 7 days prior to experimentation. For inhalation of formulations, the mice were anesthetized with isoflurane and dosed using an in-house designed micro-spray injector.

### Bioluminescence imaging

To evaluate the in vivo protein expression and tissue distribution of mFluc before and after purification, mice were administered with the same dose of mFluc-U and mFluc-P and PBS counterparts via the i.m. and i.h. routes. The Fluc expression was measured through bioluminescence imaging at 6 h, 24 h, 48 h, 72 h, and 96 h after dosing. Mice were anesthetized with isoflurane, followed by an intraperitoneal injection of the substrate D-fluorescein (3 mg/mouse). After 5 min, the bioluminescence was detected using an In Vivo Imaging System (Lumina Series III, PerkinElmer). The same dose of mFluc-U and mFluc-P and PBS counterparts were injected via i.m. and i.h. routes to evaluate the distribution of IVT mRNA transfected organs in vivo (Supplementary Fig. 2).

### Bone marrow derived dendritic cells (BMDC) maturation assay

BMDCs were harvested from the femurs of female BALB/c mice and cultured in RPMI 1640 complete medium, supplemented with 10% FBS, 1% penicillin/streptomycin, and 10 ng/mL each of IL-4 (214-14, PEPROTECH) and Granulocyte-Macrophage Colony Stimulating Factor (GM-CSF) (315-03, PEPROTECH). Fresh media replacements were performed at 2 days and 5 days post culture to discard non-adherent and weakly adherent cells, and the cultures were continued for an additional 2 days. For assessing BMDC maturation in vitro, 1×10^6^/mL BMDCs were separately co-cultured with mRBD-U, mRBD-P and PBS for 24 h. Flow cytometry staining was then conducted using Fixable Viability Stain 700 (564997, BD Biosciences), FITC anti-mouse CD11c (11-0114-81, Invitrogen), PerCP/Cy5.5 anti-mouse CD80 (104722, Biolegend), APC anti-mouse CD86 (17-0862-81, Invitrogen), and MHC Class II (I-A/I-E) Monoclonal Antibody (M5/114.15.2), eFluor™ 450, (48-5321-82, Invitrogen). Cells were incubated in Fluorescence Activated Cell Sorter (FACS) buffer for 30 min at 4°C, washed, and subsequently analyzed on a BD FACS Array flow cytometer (BD Biosciences, USA). The detailed gating strategy is shown in the Supplementary Fig. 9.

### Measurement of cytokine levels

To quantify the levels of interferon-α (IFN-α), interferon-β (IFN-β) and interleukin-6 (IL-6), DC2.4 cells were seeded at a density of 2.5×10^5^ cells/well in a 96-well plate and incubated with 100 µL of mRNA formulations at a concentration of 4 µg/mL. After 24 h of incubation, the cells were centrifuged at 1000 rpm for 5 min, and the supernatant was collected for cytokine analysis using a mouse cytokine quantification ELISA Test Kit (RX203124M for IFN-α, RX203106M for IFN-β, and RX203049M for IL-6, ruixinbio), following the manufacturer’s instructions. Subsequently, the lungs of mice administered with mRBD via intranasal (i.h.) route were collected 24 h post-administration, weighed, and homogenized in PBS at a 1:9 (m/v) ratio. The homogenates were then centrifuged at 10,000 rpm for 10 min at 4°C, and the supernatant was assayed for IFN-α, IFN-β, and IL-6 using the same ELISA kits as described for the cell culture experiments.

### Histopathology analysis

For histologic examination, major organs of mice including the heart, the liver, the spleen, the lung and the kidney, were collected and fixed in neutral 10% formalin, embedded in paraffin, sectioned and stained with hematoxylin and eosin. The sections were observed at 200-fold magnification via light microscopy (ECLIPSE 80i, Nikon). To evaluate the biochemical parameters in mouse serum, blood samples were collected from the tail vein of BALB/c mice 24 h post-administration. The serum was separated by centrifuging the collected blood at 3,000 rpm for 15 min at 4°C. The serum was then stored at −20°C until analysis. Biochemical analyses were performed by Biossci (Hubei) biotechnologies Co. Ltd., in which Serum levels of various enzymes including aspartate aminotransferase (AST) and alanine aminotransferase (ALT).

### Enzyme linked immunosorbent assay (ELISA)

Unless otherwise specified, BALB/c mice received 3 μg/mouse mRNA encapsulated in LNPs via i.m. or i.h. route. To measure the antigen-specific Immunoglobulin G (IgG) levels in serum samples and the antigen-specific secretory immunoglobulin A (sIgA) levels in bronchoalveolar lavage fluid (BALF) and nasal lavage fluid (NLF) samples, an ELISA assay was conducted. Briefly, polystyrene microtiter 96-well plates were pre-coated with 100 µL/well of RBD protein at a concentration of 2 mg/L and incubated overnight at 4°C. The plates were washed three times with PBST (PBS with 0.05% Tween-20), and blocked with 1% BSA in PBS for 1 h at 37°C to prevent non-specific binding. 100 µL/well of serial two-fold diluted serum from each mouse group was added into the pre-coated plates and incubated for 1 h at 37°C to allow for antibody binding. Detection of bound antibodies was conducted using goat anti-mouse IgG-HRP conjugate (ab97023, Abcam) or goat anti-mouse IgA-HRP conjugate (ab97235, Abcam). 100 µL of tetramethylbenzidine (TMB) substrate (ab171522, Abcam) was added to each well. The reaction was stopped after sufficient colour development, and the absorbance at 450 nm (OD_450_) was measured using a microplate reader. A well was considered positive if its OD_450_ was at least 2.1 times higher than that of the negative control which was only added PBS.

### Pseudovirus neutralization assay

The pseudotyped virus-based neutralization assay was performed as described previously(*45*). Mouse serum samples from vaccinated mice were serially diluted double-fold starting at a 1:100 dilution with DMEM containing 2% FBS for the assay. Serum was incubated with 10 μl of Luc-SARS-Cov-2 pseudotyped virus (LV-2058, PackGene, China) for 60 min, then added to the HEK293T cells stably expressing ACE2 to incubate in a standard incubator (37 °C, 5% CO_2_) for 72 h. Post infection, cells were lysed and detected using the Firefly Luciferase Reporter Gene Assay Kit. The sVNT_50_ was calculated as the serum dilution at which relative light unit (RLU) was reduced by 50% compared with RLU in virus control wells after subtraction of background RLU in cell control wells.

### Enzyme linked immunospot (ELISpot) assay

After euthanizing the mice, 10 ml of PBS was used for cardiac perfusion to remove residual circulating immune cells from the lungs. Then lungs and spleens of mice were collected at 24 h post-prime. The lungs were cut into scraps and digested with collagenase II (0.5 mg/mL, C8150, Solarbio, China) and DNase I (10 mg/ml, B002138-0100, Diamond) in CaCl_2_ (1 mM) and MgCl_2_ (1 mM) solution at 37 °C for 1 h with shaking at 220 rpm, then passed through a 40 μm cell strainer to obtain single cell suspension. The pulmonary lymphocytes were obtained by density gradient centrifugation with Percoll (17-0891-09, GE Healthcare, USA) according to the manufacturer’s instruction. In terms of splenocyte preparation, the excised spleens were ground in PBS and passed through a 40 μm cell strainer to obtain a single-cell suspension. Cell pellets were re-suspended in 3 mL of red blood cell lysis buffer (RT122-02, TIANGEN, China) for 5 min at RT to remove the red blood cells. PBS was added to wash the cells twice, then centrifuged at 1000 rpm for 5 min, the cell pellets were eventually re-suspended in RPMI1640 media supplemented with 10% FBS and 1% penicillin/streptomycin. Mouse IFN-γ/IL-4/IL-17A ELISpot PLUS kits (3321-4AST-2/3311-4APW-2/3521-4HPW-2, MabTech) were applied to measure antigen-specific IFN-γ, IL-4 and IL-17A producing cells, respectively. In brief, the single-cell suspensions were added to the plate at a concentration of 0.5×10^6^ −1.0×10^6^ cells/well. The cells were cultured with 5 μg/well SARS-CoV-2 RBD peptide pool for 48 h for 6 h at 37 °C, 5% CO_2_, the plates were washed with PBS and incubated with biotinylated anti-mouse IFN-γ, IL-4 or IL-17A antibody for 2 h at RT. Finally, TMB substrate solution was added to visualize the spots. The number of spot-forming cells was calculated on an ELISpot Reader (AID). PMA/Ionomycin was added as a positive control and PBS was used as a negative control.

### Cell staining for flow cytometry

Obtain pulmonary lymphocytes or splenocytes using the same extraction method as mentioned in the ELISpot section. The cells were initially cultured overnight in complete RPMI-1640 medium supplemented with 10% FBS and 1% penicillin/streptomycin and the SARS-CoV-2 RBD peptide pool at 5 μg/well. The cells were washed twice with PBS and adjusted to 1 × 10^6^ cells/well in flow cytometry buffer containing 0.5% FBS. For intracellular cytokine staining, cells underwent fixation and permeabilization using the Cytofix/Cytoperm kit (BD Biosciences) following the manufacturer’s guidelines. This process included a 30-min fixation at 4°C, followed by a washing step using the 1×Perm/Wash buffer provided in the kit. Subsequently, lymphocytes were stained with PE anti-mouse IFN-γ XMG1.2 (12-7311-82, Thermo Fisher Scientific) for 30 min at 4°C in the dark. Afterwards, cells were washed twice with 1×Perm/Wash buffer (421002, Biolegend) and were stained for cell surface markers. This involved incubating the cells with FITC Hamster Anti-Mouse CD3e (553061, BD Biosciences), PerCP-Cy™5.5 Rat Anti-Mouse CD4 (550954, BD Biosciences), and PE/Cyanine7 anti-mouse CD8a (100722, Biolegend) antibodies, for 30 min at 4°C in the dark, then the cells were washed twice with FACS buffer to remove unbound antibodies. Finally, the fully stained cell samples were analyzed using a flow cytometer. The pulmonary lymphocytes or splenocytes of mice were also stained at 4°C to identify effector memory T (Tem) cells, central memory T(Tcm) cells, and Trm cells, with indicated surface markers: APC anti-mouse CD44 (103012, Biolegend) and PE anti-mouse CD62L (161204, Biolegend) for Tem and Tcm cells, BV421 anti-mouse CD69 (104527, Biolegend)and BV510 anti-mouse CD103 (121423, Biolegend) for Trm cells, the cells were washed twice after staining and analyzed on a BD FACS Array flow cytometer. Data are analyzed with FlowJo software V10. The detailed gating strategy is shown in the supplementary Fig. 10-12.

### RNA sequencing

Lung tissues were collected 24 h post-prime with mRBD-U/LNP and mRBD-P/LNP via i.h. route. Total RNA was extracted using TRIzol^®^ Reagent according to the manufacturer’s protocol. Subsequent RNA purification, reverse transcription, library construction, and sequencing were conducted at Shanghai Majorbio Bio-pharm Biotechnology Co. Ltd. (Shanghai, China), following standard Illumina procedures (San Diego, CA). Gene expression levels were calculated using the Transcripts Per Million (TPM) method, and RSEM (*50*) was employed to quantify gene abundances. Differential expression analysis was conducted using DESeq2(*51*), identifying genes as differentially expressed if they exhibited |log2FC| ≥ 1 and an FDR ≤ 0.05. Functional enrichment analyses for GO and KEGG pathways were performed to determine significant enrichment of differentially expressed genes, using Goatools and KOBAS(*52*), respectively, with significance set at a Bonferroni-corrected *P*-value ≤ 0.05.

### Statistical Analysis

Unless otherwise specified, the data for all bar charts are presented as the mean ± Standard Error of Mean (SEM). Significance was determined using two-tailed unpaired *t*-test and one-way ANOVA followed by Dunnett’s multiple comparison test when the data exhibited normal distribution and variance homogeneity. GraphPad Prism 8.0 was used for data analysis. Differences were considered statistically significant when *P* < 0.05.

## ACKNOWLEDGEMENTS

This work was supported by the National Natural Science Foundation of China (NSFC, Grant No. 82173764 and 32370993), the major project of Study on Pathogenesis and Epidemic Prevention Technology System (2021YFC2302500) by the Ministry of Science and Technology of China, the Bundesministerium für Bildung und Forschung (BMBF, Grant No. 03VP10060, Zell-Trans), and the Natural Science Foundation of Chongqing (cstc2021jcyj-msxmX0136). We thank Yajuan Zhang (Southwest Hospital) for providing flow cytometry support, we also thank Dr. Xi Zhu (Shanghai Vitalgen BioPharma Co., Ltd) and Dr. Xing Liu (Sichuan ShuoGen Tech Co., Ltd) for assistance in microfluidic device.

## AUTHOR CONTRIBUTIONS

S.G. conceived, directed the project and contributed experimental materials. J.Z., Y.L., C.L., R.L., L.X., Q.X., M.L., Y.C. and X.D. performed experiments. J.Z., Y.L., D.H., C.L., S.P. and J.R. analyzed data. H.Z. and Y.L. provided the dsRNA standard. T.C. and J.Z. were responsible for the bioinformatics and RNA sequencing data analysis. S. G., X. D. and S. P. supervised the research. J.Z., C.L., Y.L. and S.G. wrote the manuscripts with help and comments from all authors.

## DATA AVAILABILITY

All data that support the findings of this study are provided within the paper and its Supplementary Information. Source data are provided with this paper.

