## Supplementary information for "Airway applied IVT mRNA vaccine needs specific sequence design and high standard purification that removes devastating dsRNA contaminant"

**This file includes:**

Supplementary Table 1 to 6

Supplementary Fig. 1 to 12

**Supplementary Table 1.** 20 genes selected from the human protein atlas (HPA) according to normalized Transcripts Per Million (nTPM) parameter.

| **Alveolar cells type 1** | **Alveolar cells type 2** | **B-cells** | **Ciliated cells** | **Club cells** | **Endothelial cells** | **Fibroblasts** | **Granulocytes** | **Macrophages** | **Smooth muscle cells** | **T-cells** |
|---|---|---|---|---|---|---|---|---|---|---|
| ACTB | FTL | CD74 | PRSS3 | FTL | TMSB10 | FTL | FTL | FTL | ACTB | ACTB |
| GPRC5A | ACTB | RPL18 | RPL18 | RPL18 | ACTB | ACTB | VIM | ACTB | VIM | TMSB10 |
| FTL | RPS5 | FTL | FTL | LYZ | FTL | DCN | RPL18 | VIM | FTL | FTL |
| GSTP1 | HSP90AA1 | RPS5 | RPS5 | GSTP1 | HSP90AA1 | VIM | ACTB | TMSB10 | RPL18 | HSP90AA1 |
| HSP90AA1 | RPL18 | ACTB | RPL6 | ACTB | VIM | TMSB10 | RPS5 | CD74 | RPS5 | RPL18 |
| CEACAM6 | GSTP1 | TMSB10 | RPLP0 | RPS5 | CD59 | RPL18 | RPL6 | HSP90AA1 | RPL6 | IL32 |
| RPL18 | RPS20 | RPL6 | GNAS | SERPINB3 | RPL18 | RPS5 | TMSB10 | CYBA | HSP90AA1 | CYBA |
| TMSB10 | RPLP0 | RPS20 | HSP90AA1 | RPL6 | TFPI | FBLN1 | RPS20 | TYROBP | TMSB10 | CST7 |
| ENO1 | RPL6 | CYBA | RPS20 | RPLP0 | RPS5 | RPL6 | HSP90AA1 | MARCO | KLF6 | RPL6 |
| RPS5 | ENO1 | RPLP0 | ACTB | TMSB10 | YBX3 | RPLP0 | LAPTM4A | RPL18 | YBX3 | RPS5 |
| MGST1 | MGST1 | HSP90AA1 | PABPC1 | RPS20 | PPP1R15A | HSP90AA1 | CAPG | LYZ | GNAS | PTPRC |
| TRAM1 | TMSB10 | VIM | SLC4A4 | HSP90AA1 | RPL6 | RPS20 | CD9 | GRN | GSTP1 | VIM |
| CD9 | CXCL2 | GNAS | VIM | CD9 | HYAL2 | GSTP1 | PEBP1 | YBX1 | MCAM | RPS20 |
| MGLL | YBX1 | PABPC1 | PEBP1 | MGST1 | KLF6 | YBX1 | RGS1 | RPS20 | CCN5 | KLF6 |
| RPL6 | KLF6 | HERPUD1 | IL32 | KLF6 | CXCL2 | RPL31 | TYROBP | RPL6 | DYNLL1 | TYROBP |
| RPS20 | GNAS | RPL31 | TMSB10 | CD74 | FKBP1A | YBX3 | CD44 | KLF6 | RPS20 | CD99 |

| | | | | | | | | | | |
|---|---|---|---|---|---|---|---|---|---|---|
| RPLP0 | RPL31 | KLF6 | NUCB2 | GNAS | RPS20 | PPP1R15A | YBX1 | RPS5 | DCN | RPLP0 |
| APLP2 | SLC25A5 | YBX1 | SLC25A5 | PABPC1 | RPLP0 | DYNLL1 | PLAUR | TYMP | RPLP0 | RPL31 |
| KLF6 | CD9 | ERP29 | RPL31 | CXCL2 | SLC9A3R2 | GNAS | CYBA | GSTP1 | YBX1 | PPP2R5C |
| CLDN18 | TRAM1 | CD79B | GSTP1 | YBX1 | YBX1 | ENO1 | RPL31 | ENO1 | LAPTM4A | CDC42 |

**Supplementary Table 2.** The 5'UTR and 3'UTR sequences of IVT mRNA applied in this study

| Gene name | 5’UTR sequence | 3’UTR sequence |
|---|---|---|
| **VIM** | GGGAGGCCCACGTATGGCGCCTCTCCAAAGG<br>CTGCAGAAGTTTCTTGCTAACAAAAAGTCCG<br>CACATTCGAGCAAAGACAGGCTTTAGCGAGT<br>TATTAAAAACTTAGGGGCGCTCTTGTCCCCCA<br>CAGGGCCCGACCGCACACAGCAAGGCGATG<br>GCCCAGCTGTAAGTTGGTAGCACTGAGAACT<br>AGCAGCGCGCGCGGAGCCCGCTGAGACTTGA<br>ATCAATCTGGTCTAACGGTTTCCCCTAAACCG<br>CTAGGAGCCCTCAATCGGCGGGACAGCAGGG<br>CGCGTCCTCTGCCACTCTCGCTCCGAGGTCCC<br>CGCGCCAGAGACGCAGCCGCGCTCCCACCAC<br>CCACACCCACCGCGCCCTCGTTCGCCTCTTCT<br>CCGGGAGCCAGTCCGCGCCACCGCCGCCGCC<br>CAGGCCATCGCCACCCTCCGCAGCCGCCACC | AAATTGCACACACTCAGTGCAGCAATATATTACCAGC<br>AAGAATAAAAAAGAAATCCATATCTTAAAGAAACAGC<br>TTTCAAGTGCCTTTCTGCAGTTTTTCAGGAGCGCAAGA<br>TAGATTTGGAATAGGAATAAGCTCTAGTTCTTAACAAC<br>CGACACTCCTACAAGATTTAGAAAAAAGTTTACAACA<br>TAATCTAGTTTACAGAAAAATCTTGTGCTAGAATACTT<br>TTTAAAAGGTATTTTGAATACCATTAAAACTGCTTTTTT<br>TTTTCCAGCAAGTATCCAACCAACTTGGTTCTGCTTCA<br>ATAAATCTTTGGAAAAACTC |
| **RPLP0** | CCTTCTCTCGCCAGGCGTCCTCGTGGAAGTGA<br>CATCGTCTTTAAACCCTGCGTGGCAATCCCTG<br>ACGCACCGCCGTGGCCACC | TCACCAAAAAGCAACCAACTTAGCCAGTTTTATTTGCA<br>AAACAAGGAAATAAAGGCTTACTTCTTTAAAAAGTC |
| **RPS20** | CTTTTTGAGGAAGACGCGGTCGTAAGGGCTG<br>AGGATTTTTGGTCCGCACGCTCCTGCTCCTGA<br>CTCACCGCTGTTCGCTCTCGCCGAGGAACAA<br>GTCGGTCAGGAAGCCCGCGCGCAACAGCCGC<br>CACC | GTCAACTATTTTAATAAATTGATGACCAGTTGTTAA |
| **RPL6** | CTCTTTCCCATCTTGCAAGGCCACC | ATGTCTTAAGAACCTAATTAAATAGCTGACTACAT |
| **RPS5** | CTCTTCCTGTCTGTACCAGGGCGGCGCGTGGT<br>CTACGCCGAGTGACAGAGACGCTCAGGCTGT<br>GTTCTCAGGGCCACC | TTTTCCCAGCTGCTGCCCAATAAACCTGTCTGCCCTTTG<br>GGGCAGTCCCAGCCA |
| **TMSB10** | GTTTCTTGCTGCAGCAACGCGAGTGGGAGCA<br>CCAGGATCTCGGGCTCGGAACGAGACTGCAC<br>GGATTGTTTTAAGAAAGCCACC | GATCCTGGAGGATTTCCTACCCCCGTCCTCTTCGAGAC<br>CCCAGTCGTGATGTGGAGGAAGAGCCACCTGCAAGAT<br>GGACACGAGCCACAAGCTGCACTGTGAACCTGGGCAC<br>TCCGCGCCGATGCCACCGGCCTGTGGGTCTCTGAAGGG<br>ACCCCCCCCCAATCGGACTGCCAAATTCTCCGGTTTGC<br>CCCGGGATATTATAGAAAATTATTTGTATGAATAATGA<br>AAATAAAACACACCTCGTGGCA |
| **RPL18** | CTCTTTCCGGACCTGGCCGAGCAGGAGGCGC<br>CATCGCCACC | CCCTGGATCCTACTCTCTTATTAAAAAGATTTTTGCTG<br>ACA |
| **HSP90AA1** | GACTGCGCAGGCGTGCTCACCTGGCGTGCTC<br>CACCCGACTGGGCGTCCGCAGGCTCCTCCCC<br>CGGGTGTGGCCTCCGGGCGGCATGGCTGCTT<br>CCCAGGTGATGCCGGCTTCAGCTAGTGGGGT<br>CTAGTTGACCGTTCCGCAGCCGCCAGGGCCA<br>GCGGAAAGCCGGTCAGGGGGAACCGCGGCG<br>GGGCTGGTGTCATGAGCCTGAGGTGAACTTG<br>AGGGTGCCTCCTCAGCGGTCTCCCGCCCTGCC<br>CTGAGGGGCGCCGGGACCCCAAAGAGCGGA<br>GGAAGAGCGCCACCCCGACGGCCACCGCTTC<br>GGAGCCAGCACGCGGGGTACCCTACGGGGA<br>GCGCGGGCCACC | TCTCTGGCTGAGGGATGACTTACCTGTTCAGTACTCTA<br>CAATTCCTCTGATAATATATTTTCAAGGATGTTTTTCTT<br>TATTTTTGTTAATATTAAAAAGTCTGTATGGCATGACA<br>ACTACTTTAAGGGGAAGATAAGATTTCTGTCTACTAAG<br>TGATGCTGTGATACCTTAGGCACTAAAGCAGAGCTAGT<br>AATGCTTTTTGAGTTTCATGTTGGTTTATTTTCACAGAT<br>TGGGGTAACGTGCACTGTAAGACGTATGTAACATGAT<br>GTTAACTTTGTGGTCTAAAGTGTTTAGCTGTCAAGCCG<br>GATGCCTAAGTAGACCAAATCTTGTTATTGAAGTGTTC<br>TGAGCTGTATCTTGATGTTTAGAAAAGTATTCGTTACA<br>TCTTGTAGGATCTACTTTTTGAACTTTTCATTCCCTGTA<br>GTTGACAATTCTGCATGTACTAGTCCTCTAGAAATAGG<br>TTAAACTGAAGCAACTTGATGGAAGGATCTCTCCACA<br>GGGCTTGTTTTCCAAAGAAAAGTATTGTTTGGAGGAGC<br>AAAGTTAAAAGCCTACCTAAGCATATCGTAAAGCTGTT |

| | | |
|---|---|---|
| | | CAAAAATAACTCAGACCCAGTCTTGTGGATGGAAATG<br>TAGTGCTCGAGTCACATTCTGCTTAAAGTTGTAACAAA<br>TACAGATGAGTTAAAAGATATTGTGTGACAGTGTCTTA<br>TTTAGGGGGAAAGGGGAGTATCTGGATGACAGTTAGT<br>GCCAAAATGTAAAACATGAGGCGCTAGCAGGAGATGG<br>TTAAACACTAGCTGCTCCAAGGGTTGACATGGTCTTCC<br>CAGCATGTACTCAGCAGGTGTGGGGTGGAGCACACGT<br>AGGCACAGAAAACAGGAATGCAGACAACATGCATCCC<br>CTGCGTCCATGAGTTACATGTGTTCTCTTAGTGTCCAC<br>GTTGTTTTGATGTTATTCATGGAATACCTTCTGTGTTAA<br>ATACAGTCACTTAATTCCTTGGC |
| **FTL** | GCAGTTCGGCGGTCCCGCGGGTCTGTCTCTTG<br>CTTCAACAGTGTTTGGACGGAACAGATCCGG<br>GGACTCTCTTCCAGCCTCCGACCGCCCTCCGA<br>TTTCCTCTCCGCTTGCAACCTCCGGGACCATC<br>TTCTCGGCCATCTCCTGCTTCTGGGACCTGCC<br>AGCACCGTTTTTGTGGTTAGCTCCTTCTTGCC<br>AACCAACCGCCACC | GAGCCTTCTGAGCCCAGCGACTTCTGAAGGGCCCCTTG<br>CAAAGTAATAGGGCTTCTGCCTAAGCCTCTCCCTCCAG<br>CCAATAGGCAGCTTTCTTAACTATCCTAACAAGCCTTG<br>GACCAAATGGAAATAAAGCTTTTTGATGCA |
| **ACTB** | ACCGCCGAGACCGCGTCCGCCCCGCGAGCAC<br>AGAGCCTCGCCTTTGCCGATCCGCCGCCCGTC<br>CACACCCGCCGCCAGCTCACCGCCACC | GCGGACTATGACTTAGTTGCGTTACACCCTTTCTTGAC<br>AAAACCTAACTTGCGCAGAAAACAAGATGAGATTGGC<br>ATGGCTTTATTTGTTTTTTTTGTTTTGTTTTGGTTTTTTT<br>TTTTTTTTTGGCTTGACTCAGGATTTAAAAACTGGAAC<br>GGTGAAGGTGACAGCAGTCGGTTGGAGCGAGCATCCC<br>CCAAAGTTCACAATGTGGCCGAGGACTTTGATTGCACA<br>TTGTTGTTTTTTTAATAGTCATTCCAAATATGAGATGCG<br>TTGTTACAGGAAGTCCCTTGCCATCCTAAAAGCCACCC<br>CACTTCTCTCTAAGGAGAATGGCCCAGTCCTCTCCCAA<br>GTCCACACAGGGGAGGTGATAGCATTGCTTTCGTGTAA<br>ATTATGTAATGCAAAATTTTTTTAATCTTCGCCTTAATA<br>CTTTTTTATTTTGTTTTATTTTGAATGATGAGCCTTCGT<br>GCCCCCCCTTCCCCCTTTTTTGTCCCCCAACTTGAGATG<br>TATGAAGGCTTTTGGTCTCCCTGGGAGTGGGTGGAGGC<br>AGCCAGGGCTTACCTGTACACTGACTTGAGACCAGTTG<br>AATAAAAGTGCACACCTTAAAAATGA |
| **Moderna (mRNA-1273)** | GGGAAATAAGAGAGAAAAGAAGAGTAAGAA<br>GAAATATAAGACCCCGGCGCCGCCACC | GCTGGAGCCTCGGTGGCCTAGCTTCTTGCCCCTTGGGC<br>CTCCCCCCAGCCCCTCCTCCCCTTCCTGCACCCGTACCC<br>CCGTGGTCTTTGAATAAAGTCTGAGTGGGCGGCA |
| **BioNTech (BNT162b2)** | GAGAATAAACTAGTATTCTTCTGGTCCCCAC<br>AGACTCAGAGAGAACCCGCCACC | CTCGAGCTGGTACTGCATGCACGCAATGCTAGCTGCCC<br>CTTTCCCGTCCTGGGTACCCCGAGTCTCCCCCGACCTC<br>GGGTCCCAGGTATGCTCCCACCTCCACCTGCCCCACTC<br>ACCACCTCTGCTAGTTCCAGACACCTCCCAAGCACGCA<br>GCAATGCAGCTCAAAACGCTTAGCCTAGCCACACCCC<br>CACGGGAAACAGCAGTGATTAACCTTTAGCAATAAAC<br>GAAAGTTTAACTAAGCTATACTAACCCCAGGGTTGGTC<br>AATTTCGTGCCAGCCACACCCTGGAGCTAGC |
| **TB-Opti** | GTTTCTTGCTGCAGCAACGCGAGTGGGAGCA<br>CCAGGATCTCGGGCTCGGAACGAGACTGCAC<br>GGATTGTTTTAAGAAAGCCACC | GATCCTGGAGGATTTCCTCCTCTTCGAGTCGCCGGTCG<br>GTTCTCCGTAAATCGTGGCA |

The underlined part in the 5'UTR sequence represents the Kozak sequence.

**Supplementary Table 3.** The open reading frame (ORF) sequences of IVT mRNA adopted in this study

| mRNA name | sequence |
|---|---|
| **mFLUC** | ATGGAAGATGCCAAAAACATTAAGAAGGGCCCAGCGCCATTCTACCCACTCGAAGACGGGACCGCCGGC<br>GAGCAGCTGCACAAAGCCATGAAGCGCTACGCCCTGGTGCCCGGCACCATCGCCTTTACCGACGCACATA<br>TCGAGGTGGACATTACCTACGCCGAGTACTTCGAGATGAGCGTTCGGCTGGCAGAAGCTATGAAGCGCTA<br>TGGGCTGAATACAAACCATCGGATCGTGGTGTGCAGCGAGAATAGCTTGCAGTTCTTCATGCCCGTGTTG<br>GGTGCCCTGTTCATCGGTGTGGCTGTGGCCCCAGCTAACGACATCTACAACGAGCGCGAGCTGCTGAACA<br>GCATGGGCATCAGCCAGCCCACCGTCGTATTCGTGAGCAAGAAAGGGCTGCAAAAGATCCTCAACGTGCA<br>AAAGAAGCTACCGATCATACAAAAGATCATCATCATGGATAGCAAGACCGACTACCAGGGCTTCCAAAG<br>CATGTACACCTTCGTGACTTCCCATTTGCCACCCGGCTTCAACGAGTACGACTTCGTGCCCGAGAGCTTCG<br>ACCGGGACAAAACCATCGCCCTGATCATGAACAGTAGTGGCAGTACCGGATTGCCCAAGGGCGTAGCCCT<br>ACCGCACCGCACCGCTTGTGTCCGATTCAGTCATGCCCGCGACCCCATCTTCGGCAACCAGATCATCCCCG<br>ACACCGCTATCCTCAGCGTGGTGCCATTTCACCACGGCTTCGGCATGTTCACCACGCTGGGCTACTTGATC<br>TGCGGCTTTCGGGTCGTGCTCATGTACCGCTTCGAGGAGGAGCTATTCTTGCGCAGCTTGCAAGACTATAA<br>GATTCAATCTGCCCTGCTGGTGCCCACACTATTTAGCTTCTTCGCTAAGAGCACTCTCATCGACAAGTACG<br>ACCTAAGCAACTTGCACGAGATCGCCAGCGGCGGGGCGCCGCTCAGCAAGGAGGTAGGTGAGGCCGTGG<br>CCAAACGCTTCCACCTACCAGGCATCCGCCAGGGCTACGGCCTGACAGAAACAACCAGCGCCATTCTGAT<br>CACCCCCGAAGGGGACGACAAGCCTGGCGCAGTAGGCAAGGTGGTGCCCTTCTTCGAGGCTAAGGTGGT<br>GGACTTGGACACCGGTAAGACACTGGGTGTGAACCAGCGCGGCGAGCTGTGCGTCCGTGGCCCCATGATC<br>ATGAGCGGCTACGTTAACAACCCCGAGGCTACAAACGCTCTCATCGACAAGGACGGCTGGCTGCACAGCG<br>GCGACATCGCCTACTGGGACGAGGACGAGCACTTCTTCATCGTGGACCGGCTGAAGAGCCTGATCAAATA<br>CAAGGGCTACCAGGTAGCCCCAGCCGAACTGGAGAGCATCCTGCTGCAACACCCCAACATCTTCGACGCC<br>GGGGTCGCCGGCCTGCCCGACGACGATGCCGGCGAGCTGCCCGCCGCAGTCGTCGTGCTGGAACACGGTA<br>AAACCATGACCGAGAAGGAGATCGTGGACTATGTGGCCAGCCAGGTTACAACCGCCAAGAAGCTGCGCG<br>GTGGTGTTGTGTTCGTGGACGAGGTGCCTAAAGGACTGACCGGCAAGTTGGACGCCCGCAAGATCCGCGA<br>GATTCTCATTAAGGCCAAGAAGGGCGGCAAGATCGCCGTGTGATAATAG |
| **mEGFP** | ATGGTGAGCAAGGGCGAGGAGCTGTTCACCGGGGTGGTGCCCATCCTGGTCGAGCTGGACGGCGACGTA<br>AACGGCCACAAGTTCAGCGTGTCCGGCGAGGGCGAGGGCGATGCCACCTACGGCAAGCTGACCCTGAAG<br>TTCATCTGCACCACCGGCAAGCTGCCCGTGCCCTGGCCCACCCTCGTGACCACCCTGACCTACGGCGTGCA<br>GTGCTTCAGCCGCTACCCCGACCACATGAAGCAGCACGACTTCTTCAAGTCCGCCATGCCCGAAGGCTAC<br>GTCCAGGAGCGCACCATCTTCTTCAAGGACGACGGCAACTACAAGACCCGCGCCGAGGTGAAGTTCGAG<br>GGCGACACCCTGGTGAACCGCATCGAGCTGAAGGGCATCGACTTCAAGGAGGACGGCAACATCCTGGGG<br>CACAAGCTGGAGTACAACTACAACAGCCACAACGTCTATATCATGGCCGACAAGCAGAAGAACGGCATC<br>AAGGTGAACTTCAAGATCCGCCACAACATCGAGGACGGCAGCGTGCAGCTCGCCGACCACTACCAGCAG<br>AACACCCCCATCGGCGACGGCCCCGTGCTGCTGCCCGACAACCACTACCTGAGCACCCAGTCCGCCCTGA<br>GCAAAGACCCCAACGAGAAGCGCGATCACATGGTCCTGCTGGAGTTCGTGACCGCCGCCGGGATCACTCT<br>CGGCATGGACGAGCTGTACAAGTAA |
| **mRBD** | ATGAGAGTCCAACCAACAGAATCTATTGTTAGATTTCCTAATATTACAAACTTGTGCCCTTTTGATGAAGT<br>TTTTAACGCCACCAGATTTGCATCTGTTTATGCTTGGAACAGGAAGAGAATCAGCAACTGTGTTGCTGATT<br>ATTCTGTCCTATATAATCTCGCATCATTTTTCACTTTTAAGTGTTATGGAGTGTCTCCTACTAAATTAAATG<br>ATCTCTGCTTTACTAATGTCTATGCAGATTCATTTGTAATTAGAGGTGATGAAGTCAGACAAATCGCTCCA<br>GGGCAAACTGGAAATATTGCTGATTATAATTATAAATTACCAGATGATTTTACAGGCTGCGTTATAGCTTG<br>GAATTCTAACAAGCTTGATTCTAAGGTTAGTGGTAATTATAATTACCTGTATAGATTGTTTAGGAAGTCTA<br>ATCTCAAACCTTTTGAGAGAGATATTTCAACTGAAATCTATCAGGCCGGTAACAAACCTTGTAATGGTGTT<br>GCAGGTTTTAATTGTTACTTTCCTTTACAATCATATGGTTTCCAACCCACTAATGGTGTTGGTTACCAACCA<br>TACAGAGTAGTAGTACTTTCTTTTGAACTTCTACATGCACCAGCAACTGTTTGTGGACCTAAAAAGTAA |
| **mSpike** | ATGTTTGTTTTTCTTGTTTTATTGCCACTAGTCTCTAGTCAGTGTGTTAATCTTACAACCAGAACTCAATTA<br>CCCCCTGCATACACTAATTCTTTCACACGTGGTGTTTATTACCCTGACAAAGTTTTCAGATCCTCAGTTTTA<br>CATTCAACTCAGGACTTGTTCTTACCTTTCTTTTCCAATGTTACTTGGTTCCATGTTATCTCTGGGACCAAT<br>GGTACTAAGAGGTTTGATAACCCTGTCCTACCATTTAATGATGGTGTTTATTTTGCTTCCATTGAGAAGTCT<br>AACATAATAAGAGGCTGGATTTTTGGTACTACTTTAGATTCGAAGACCCAGTCCCTACTTATTGTTAATAA<br>CGCTACTAATGTTGTTATTAAAGTCTGTGAATTTCAATTTTGTAATGATCCATTTTTGGACCACAAAAACA<br>ACAAAAGTTGGATGGAAAGTGAGTTCAGAGTTTATTCTAGTGCGAATAATTGCACTTTTGAATATGTCTCT<br>CAGCCTTTTCTTATGGACCTTGAAGGAAAACAGGGTAATTTCAAAAATCTTAGGGAATTTGTGTTTAAGAA<br>TATTGATGGTTATTTTAAAATATATTCTAAGCACACGCCTATTATAGTGCGTGATCTCCCTCAGGGTTTTTC<br>GGCTTTAGAACCATTGGTAGATTTGCCAATAGGTATTAACATCACTAGGTTTCAAACTTTACTTGCTTTAC<br>ATAGAAGTTATTTGACTCCTGGTGATTCTTCTTCAGGTTGGACAGCTGGTGCTGCAGCTTATTATGTGGGT |

TATCTTCAACCTAGGACTTTTCTATTAAAATATAATGAAAATGGAACCATTACAGATGCTGTAGACTGTGC
ACTTGACCCTCTCTCAGAAACAAAGTGTACGTTGAAATCCTTCACTGTAGAAAAAGGAATCTATCAAACT
TCTAACTTTAGAGTCCAACCAACAGAATCTATTGTTAGATTTCCTAATATTACAAACTTGTGCCCTTTTGAT
GAAGTTTTTAACGCCACCAGATTTGCATCTGTTTATGCTTGGAACAGGAAGAGAATCAGCAACTGTGTTGC
TGATTATTCTGTCCTATATAATCTCGCATCATTTTTCACTTTTAAGTGTTATGGAGTGTCTCCTACTAAATT
AAATGATCTCTGCTTTACTAATGTCTATGCAGATTCATTTGTAATTAGAGGTGATGAAGTCAGACAAATCG
CTCCAGGGCAAACTGGAAATATTGCTGATTATAATTATAAATTACCAGATGATTTTACAGGCTGCGTTATA
GCTTGGAATTCTAACAAGCTTGATTCTAAGGTTAGTGGTAATTATAATTACCTGTATAGATTGTTTAGGAA
GTCTAATCTCAAACCTTTTGAGAGAGATATTTCAACTGAAATCTATCAGGCCGGTAACAAACCTTGTAATG
GTGTTGCAGGTTTTAATTGTTACTTTCCTTTACAATCATATGGTTTCCAACCCACTAATGGTGTTGGTTACC
AACCATACAGAGTAGTAGTACTTTCTTTTGAACTTCTACATGCACCAGCAACTGTTTGTGGACCTAAAAAG
TCTACTAATTTGGTTAAAAACAAATGTGTCAATTTCAACTTCAATGGTTTAAAAGGCACAGGTGTTCTTAC
TGAGTCTAACAAAAAGTTTCTGCCTTTCCAACAATTTGGCAGAGACATTGCTGACACTACTGATGCTGTCC
GTGATCCACAGACACTTGAGATTCTTGACATTACACCATGTTCTTTTGGTGGTGTCAGTGTTATAACACCA
GGAACAAATACTTCTAACCAGGTTGCTGTTCTTTATCAGGGTGTTAACTGCACAGAAGTCCCTGTTGCTAT
TCATGCAGATCAACTTACTCCTACTTGGCGTGTTTATTCTACAGGTTCTAATGTTTTCAAACACGTGCAGG
CTGTTTAATAGGGGCTGAATATGTCAACAACTCATATGAGTGTGACATACCCATTGGTGCAGGTATATGC
GCTAGTTATCAGACTCAGACTAAGTCTCATCGGCGGGCACGTAGTGTAGCTAGTCAATCCATCATTGCCTA
CACTATGTCACTTGGTGCAGAAAATTCAGTTGCTTACTCTAATAACTCTATTGCCATACCCACAAATTTTA
CTATTAGTGTTACCACAGAAATTCTACCAGTGTCTATGACCAAGACATCAGTAGATTGTACAATGTACATT
TGTGGTGATTCAACTGAATGCAGCAATCTTTTGTTGCAATATGGCAGTTTTTGTACACAATTAAACCGTGC
TTTAACTGGAATAGCTGTTGAACAAGACAAAAACACCCAAGAAGTTTTTGCACAAGTCAAACAAATTTAC
AAAACACCACCAATTAAATATTTTGGTGGTTTTAATTTTTCACAAATATTACCAGATCCATCAAAACCAAG
CAAGAGGTCATTTATTGAAGATCTACTTTTCAACAAAGTGACACTTGCAGATGCTGGCTTCATCAAACAAT
ATGGTGATTGCCTTGGTGATATTGCTGCTAGAGACCTCATTTGTGCACAAAAGTTTAAAGGCCTTACTGTT
TTGCCACCTTTGCTCACAGATGAAATGATTGCTCAATACACTTCTGCACTGTTAGCGGGTACAATCACTTC
TGGTTGGACCTTTGGTGCAGGTGCTGCATTACAAATACCATTTGCTATGCAAATGGCTTATAGGTTTAATG
GTATTGGAGTTACACAGAATGTTCTCTATGAGAACCAAAAATTGATTGCCAACCAATTTAATAGTGCTATT
GGCAAAATTCAAGACTCACTTTCTTCCACAGCAAGTGCACTTGGAAAACTTCAAGATGTGGTCAACCATA
ATGCACAAGCTTTAAACACGCTTGTTAAACAACTTAGCTCCAAATTTGGTGCAATTTCAAGTGTTTTAAAT
GATATCTTTTCACGTCTTGACAAAGTTGAGGCTGAAGTGCAAATTGATAGGTTGATCACAGGCAGACTTC
AAAGTTTGCAGACATATGTGACTCAACAATTAATTAGAGCTGCAGAAATCAGAGCTTCTGCTAATCTTGCT
GCTACTAAAATGTCAGAGTGTGTACTTGGACAATCAAAAAGAGTTGATTTTTGTGGAAAGGGCTATCATC
TTATGTCCTTCCCTCAGTCAGCACCTCATGGTGTAGTCTTCTTGCATGTGACTTATGTCCCTGCACAAGAAA
AGAACTTCACAACTGCTCCTGCCATTTGTCATGATGGAAAAGCACACTTTCCTCGTGAAGGTGTCTTTGTT
TCAAATGGCACACACTGGTTTGTAACACAAAGGAATTTTTATGAACCACAAATCATTACTACAGACAACA
CATTTGTGTCTGGTAACTGTGATGTTGTAATAGGAATTGTCAACAACACAGTTTATGATCCTTTGCAACCT
GAATTAGACTCATTCAAGGAGGAGTTAGATAAATATTTTAAGAATCATACATCACCAGATGTTGATTTAG
GTGACATCTCTGGCATTAATGCTTCAGTTGTAAACATTCAAAAAGAAATTGACCGCCTCAATGAGGTTGCC
AAGAATTTAAATGAATCTCTCATCGATCTCCAAGAACTTGGAAAGTATGAGCAGTATATAAAATGGCCAT
GGTACATTTGGCTAGGTTTTATAGCTGGCTTGATTGCCATAGTAATGGTGACAATTATGCTTTGCTGTATG
ACCAGTTGCTGTAGTTGTCTCAAGGGCTGTTGTTCTTGTGGATCCTGCTGCAAATTTGATGAAGACGACTC
TGAGCCAGTGCTCAAAGGAGTCAAATTACATTACACATAATGA

**Supplementary Table 4.** The IVT mRNA functional element sequences used in the sequence optimization process

| Element name | sequence |
|---|---|
| Apt-eIF4G | ACTCACTATTTGTTTTCGCGCCCAGTTGCAAAAAGTGTCG |
| R3U | AAAACTCAATGTATTTCTGAGGAAGCGTGGTGCATAATGCCACGCAGCGTC TGCATAACTTTTATTATTTCTTTTATTAATCAACAAA |
| Kozak | GCCACC |

**Supplementary Table 5.** Method for Purification of IVT mRNA by HPLC.

| Time（min） | MPA (%) | MPB (%) | MPC (%) | MPD (%) | Flow (mL/min) |
|---|---|---|---|---|---|
| 0.00 | 100 | 0 | 0 | 0 | 1 |
| 2.00 | 100 | 0 | 0 | 0 | 1 |
| 2.01 | 0 | 100 | 0 | 0 | 1 |
| 6.00 | 0 | 100 | 0 | 0 | 1 |
| 6.01 | 0 | 0 | 100 | 0 | 1 |
| 8.00 | 0 | 0 | 100 | 0 | 1 |
| 8.01 | 0 | 0 | 0 | 100 | 1 |
| 10.00 | 0 | 0 | 0 | 100 | 1 |

MPA-D: mobile phase A-D.

**Supplementary Table 6.** Peptide Pools of 14-mer Overlapping Peptides Spanning the SARS-CoV-2 RBD Protein

| NO | Sequence | NO | Sequence |
|---|---|---|---|
| 1 | RVQPTESIVRFPNI | 23 | FTGCVIAWNSNNLD |
| 2 | ESIVRFPNITNLCP | 24 | IAWNSNNLDSKVGG |
| 3 | FPNITNLCPFGEVF | 25 | NNLDSKVGGNYNYL |
| 4 | NLCPFGEVFNATRF | 26 | KVGGNYNYLYRLFR |
| 5 | GEVFNATRFASVYA | 27 | YNYLYRLFRKSNLK |
| 6 | ATRFASVYAWNRKR | 28 | RLFRKSNLKPFERD |
| 7 | SVYAWNRKRISNCV | 29 | SNLKPFERDISTEI |
| 8 | NRKRISNCVADYSV | 30 | FERDISTEIYQAGS |
| 9 | SNCVADYSVLYNSA | 31 | STEIYQAGSTPCNG |
| 10 | DYSVLYNSASFSTF | 32 | QAGSTPCNGVEGFN |
| 11 | YNSASFSTFKCYGV | 33 | PCNGVEGFNCYFPL |
| 12 | FSTFKCYGVSPTKL | 34 | EGFNCYFPLQSYGF |
| 13 | CYGVSPTKLNDLCF | 35 | YFPLQSYGFQPTNG |
| 14 | PTKLNDLCFTNVYA | 36 | SYGFQPTNGVGYQP |
| 15 | DLCFTNVYADSFVI | 37 | PTNGVGYQPYRVVV |
| 16 | NVYADSFVIRGDEV | 38 | GYQPYRVVVLSFEL |
| 17 | SFVIRGDEVRQIAP | 39 | RVVVLSFELLHAPA |
| 18 | GDEVRQIAPGQTGK | 40 | SFELLHAPATVCGP |
| 19 | QIAPGQTGKIADYN | 41 | HAPATVCGPKKSTN |
| 20 | QTGKIADYNYKLPD | 42 | VCGPKKSTNLVKNK |
| 21 | ADYNYKLPDDFTGC | 43 | KSTNLVKNKCVNF |
| 22 | KLPDDFTGCVIAWN | | |

### SUPPLEMENTARY FIGURES

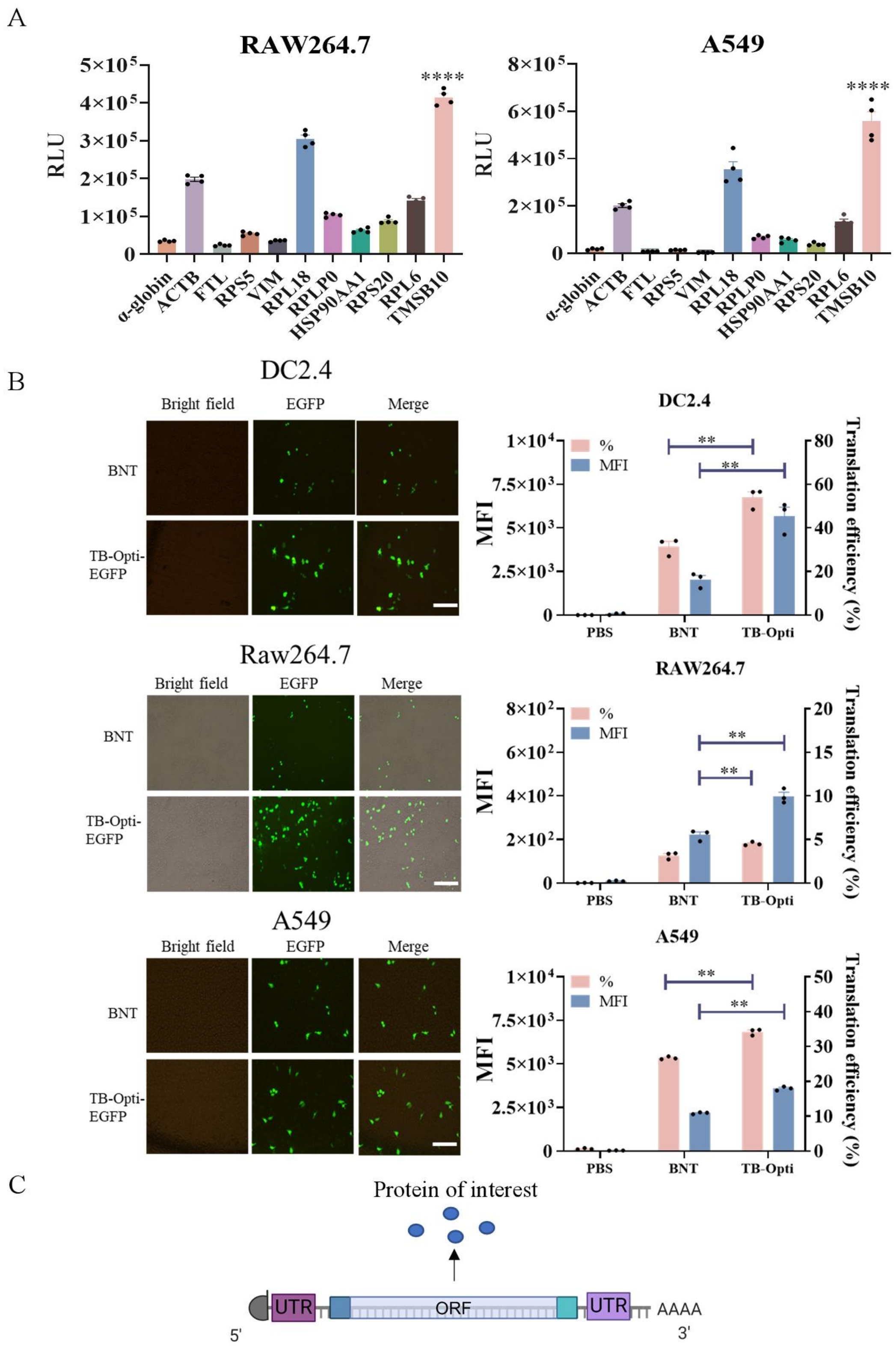


**Supplementary Fig. 1. In vitro screening and modification of UTRs of IVT mRNA sequences encoding various proteins of interest.** (A) Expression of IVT mRNA constructs designed with different 5′UTRs and 3′UTRs derived from selected genes. These constructs were synthesized and transfected into Raw264.7 cells and A549 cells to assess their impact on mRNA transfection efficiency. (B) Fluorescence microscopy images and flow cytometry analysis evaluating the expression levels of mEGFP designed with UTR of TB Opti, BioNTech BNT-162b2 (BNT), and Moderna mRNA-1273 (MDN) in various cells. (C) Schematic of the modification of mRNA encoding various proteins of interest. Statistical significance was assessed by two-tailed unpaired *t*-test and one-way ANOVA with Dunnett's post-hoc test (**$P < 0.01$; ****$P < 0.0001$). DC2.4: mouse dendritic cells; RAW264.7: mouse mononuclear macrophage leukemia cells; A549: human non-small cell lung cancer cells.

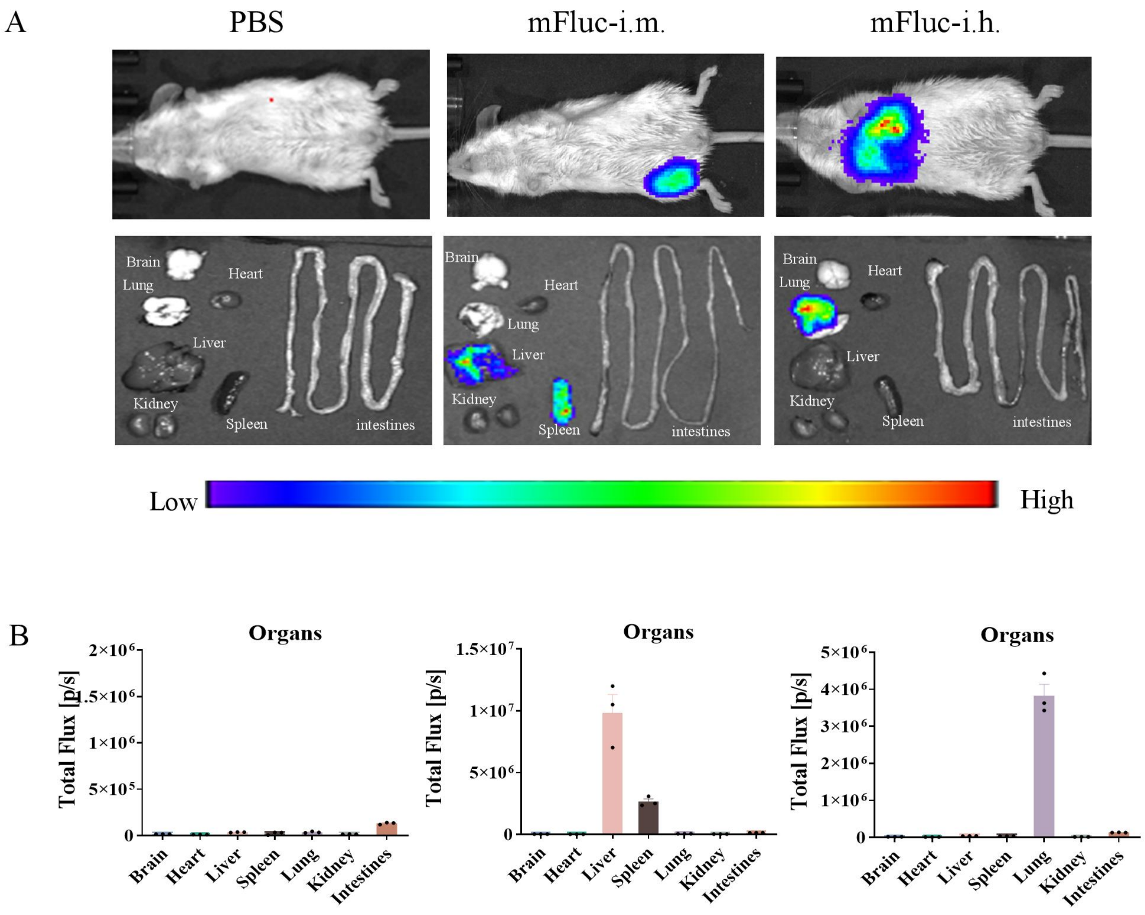


**Supplementary Fig. 2. Bioluminescence imaging of Fluc expression in mice and excised organs following i.m. or i.h. route administration.** The mFluc was designed using the TB-Opti sequence, encapsulated into LNP and administered via intramuscular (i.m.) or intratracheal (i.h.) routes. The control group received PBS via the i.h. route. Signals were detected using IVIS to show the bio-distribution of mFluc expression post-administration in mice.

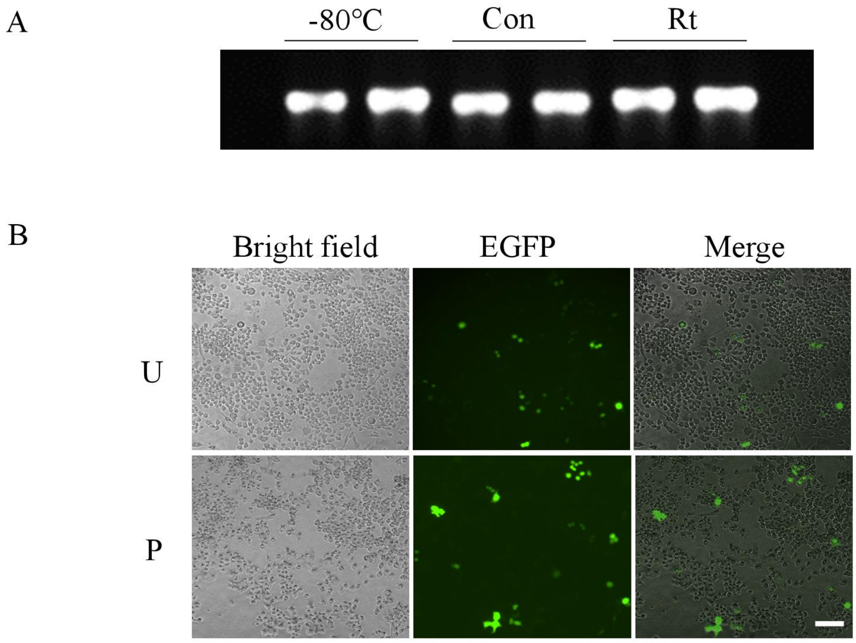


**Supplementary Fig. 3. Evaluations of the impact of vacuum concentration on purified mRNA, and the expression of purified mEGFP in DC2.4.** (A) Denatured agarose gel electrophoresis of purified mRNA concentrated at room temperature (RT) under vacuum conditions, there was no difference in the integrity with purified mRNA stewing at -80 ℃ or RT. (B) Representative transfection profiles of unpurified mEGFP (U) or purified mEGFP (P) in DC2.4 cells 24 h post-transfection. Images were obtained by fluorescence microscopy with equal acquisition parameters. Scale bar, 100 μm.

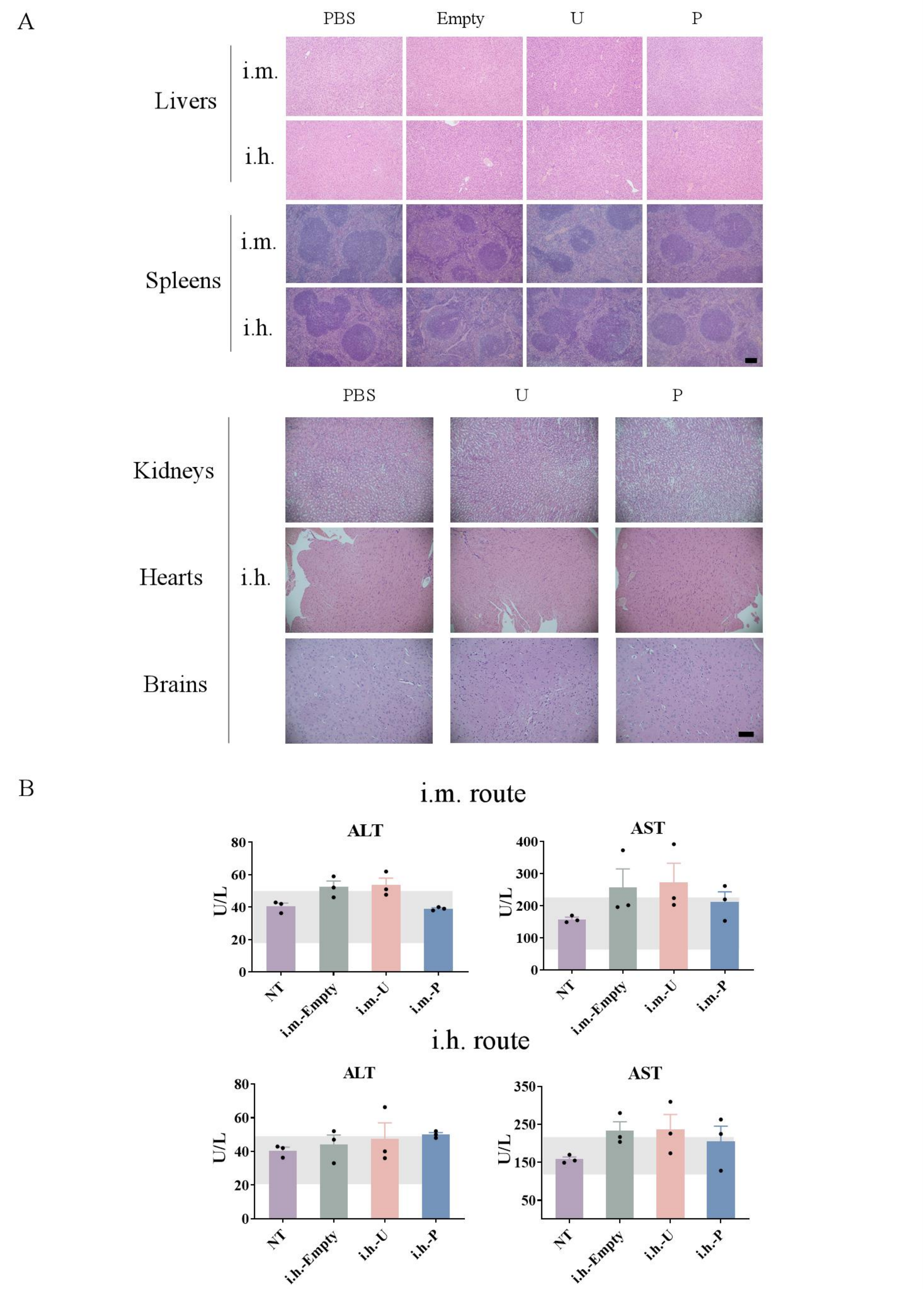


**Supplementary Fig. 4. Impacts of purification on in vivo toxicity.** (**A**) Representative hematoxylin and eosin (H&E) staining of major organs of mice collected 24 h after administration (i.m. and i.h.) of unpurified mRBD (U) and purified mRBD (P). Samples treated by PBS and LNP formulation without IVT mRNA (Empty) were used as controls. Scale bar, 100 μm. (**B**) Complete blood chemistry metrics of mice treated by Empty, mRBD-U (U) and mRBD-P (P) via i.m. or i.h. route. ALT: alanine aminotransferase, AST: aspartate aminotransferase. The grey regions represent the normal range of each parameter as a reference.

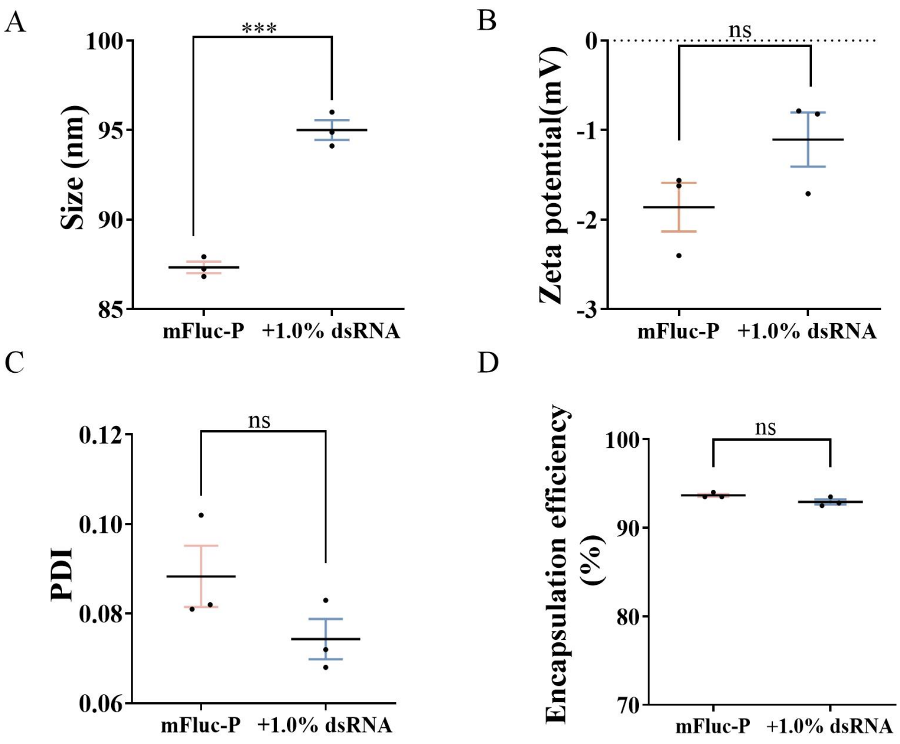


**Supplementary Fig. 5. Physicochemical characterizations of LNP encapsulated with mFluc-P or mFluc-P+dsRNA.** Dynamic light scattering measurement (hydrodynamic size (A), zeta potential (B) and polydispersity index (PDI) (C)) and encapsulation efficiency (D) of purified mFLuc (mFluc-P) or purified mFLuc with addition of 1.0% (w/w) dsRNA (mFluc-P+dsRNA). Statistical significance was assessed by two-tailed unpaired *t*-test (ns: not significant; ***$P < 0.001$).

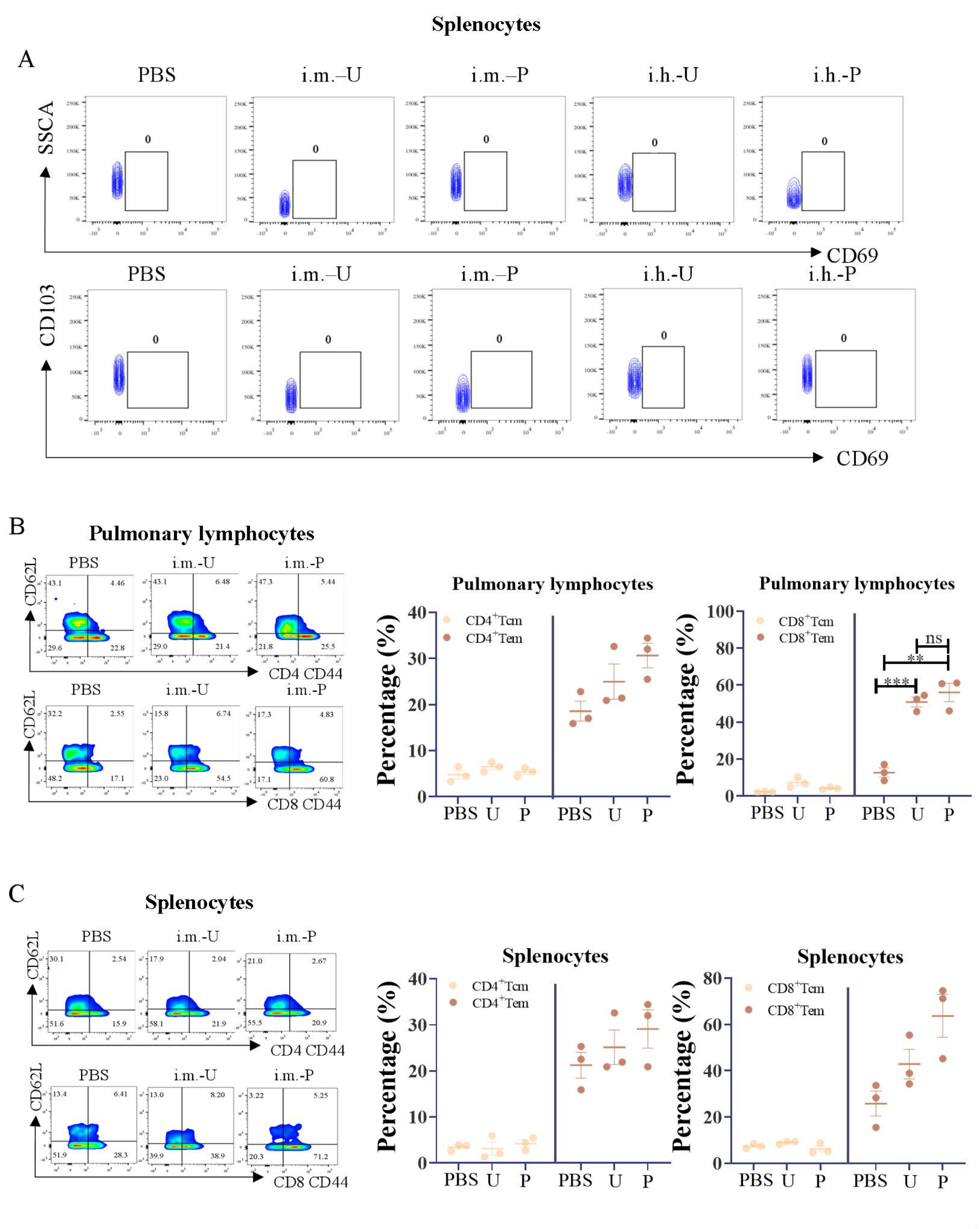


**Supplementary Fig. 6. Immunologic evaluations of SARS-CoV-2 specific T cell responses post-immunization with unpurified mRBD (U) or purified mRBD (P) via i.m. or i.h. route.** (A) Tissue-resident memory T cell (Trm) frequencies across CD4$^{+}$ (CD4$^{+}$CD44$^{+}$CD69$^{+}$) and CD8$^{+}$ (CD8$^{+}$CD44$^{+}$CD69$^{+}$CD103$^{+}$) T cells in the spleens of mice vaccinated via i.m. routes. (B, C) Levels of central memory (Tcm, CD44$^{+}$CD62L$^{+}$) and effector memory (Tem, CD44$^{+}$CD62L$^{-}$) T cells across CD4$^{+}$ and CD8$^{+}$T cells in the lungs (B) or spleens (C) of mice vaccinated via i.m. route. Each symbol in B-C represents one biologically independent animal. Statistical significance was calculated by one-way ANOVA with Dunnett's post-hoc test (ns: not significant; $^{**}P < 0.01$; $^{***}P < 0.001$).

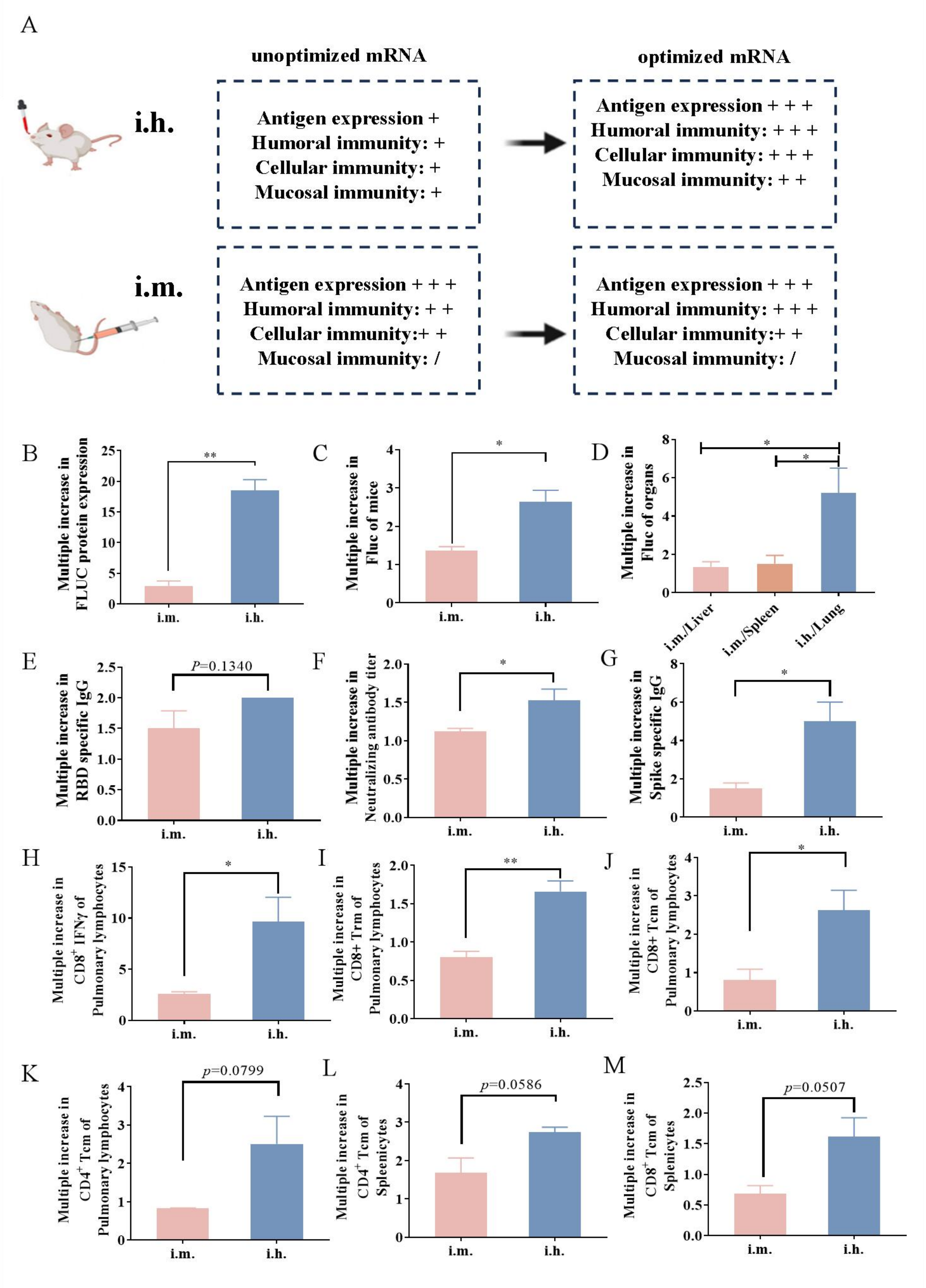


**Supplementary Fig. 7. Comparison on the increased rates of various biological parameters mediated by the optimized IVT mRNA (specific UTR design plus purification) inoculated via i.m. and i.h. routes.** (A) A summary diagram comparing the impacts of optimized vs. unoptimized mRNA vaccines administered via i.h. and i.m. routes. (B) Increased folds of total Fluc protein expression mediated by optimized mRNA compared to unoptimized counterpart in live mice 120 h post-administration by i.m. and i.h. routes. (C, D) Increased folds of Fluc protein expression in live mice (C) and primary organs (D) 6 h post-administration of optimized mRNA compared to unoptimized counterpart. (E, F) Increased folds of humoral immune response induced by mRBD-P compared to mRBD-U via i.m. and i.h. routes, including antigen-specific IgG levels (E) and pseudovirus neutralizing antibody titers (F). (G) Enhanced folds of antigen-specific IgG levels induced by mSpike-P compared to mSpike-U via i.m. and i.h. routes. (H-M) Elevated folds of cellular immune responses induced by mRBD-P compared to mRBD-U via i.m. and i.h. routes: $CD8^{+}$IFN-γ (H), $CD8^{+}$Trm (I), $CD8^{+}$Tcm (J), $CD4^{+}$Tcm (K) in lung lymphocytes and $CD8^{+}$ Tcm (M), $CD4^{+}$ Tcm (M) in splenocyte. Statistical significance was assessed by two-tailed unpaired *t*-test and one-way ANOVA with Dunnett's post-hoc test (ns: not significant; *$P < 0.05$, **$P < 0.01$).

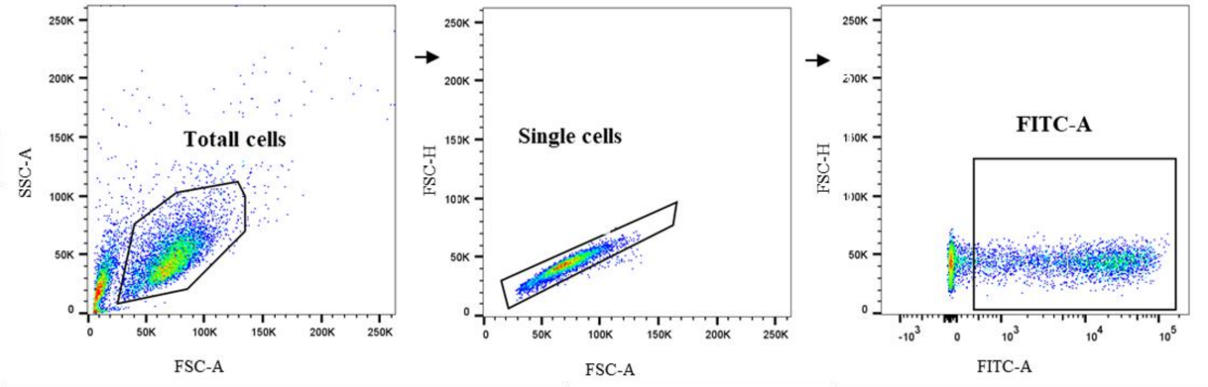


**Supplementary Fig. 8. Gating strategy for identification of cells successfully expressing EGFP.** The strategy begins by gating on cells using forward scatter-area and side scatter-area (FSC-A and SSC-A) to exclude debris. Next, gating on cells using FSC-A and side scatter height (SSC-H) to exclude adhesive cells. A fluorescence gate is applied to isolate cells exhibiting EGFP-specific fluorescence, ensuring accurate identification of cells successfully expressing EGFP.

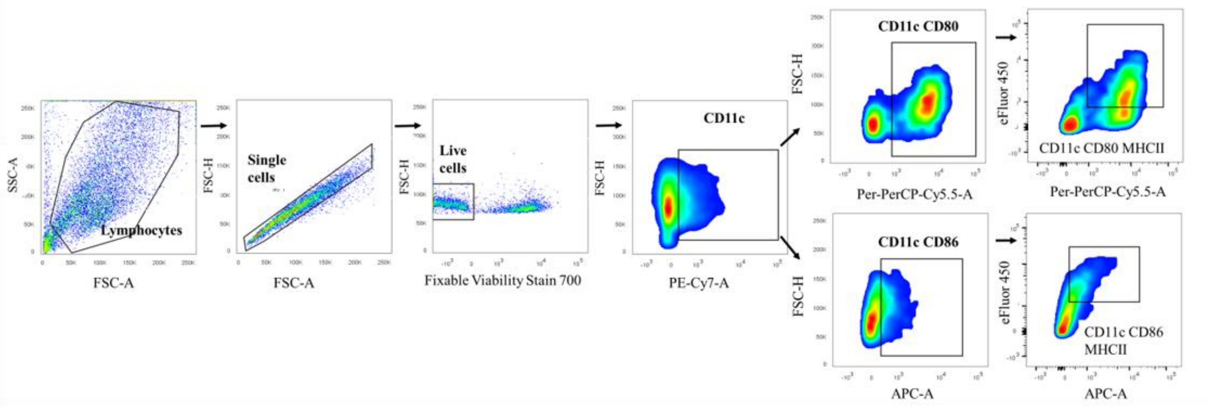


**Supplementary Fig. 9. Gating strategy for identification of expression of $CD11c^{+}CD80^{+}$ and $CD80^{+}$MHC-$II^{+}$ (gated on $CD11c^{+}$) or $CD11c^{+}CD86^{+}$and $CD86^{+}$MHC-$II^{+}$ (gated on $CD11c^{+}$) on bone marrow derived dendritic cells (BMDCs).** Initially, gating on cells based on FSC-A and SSC-A to exclude debris. Next, using FSC-A and FSC-H to exclude adhesive cells, Fixable Viability Stain 700 to exclude dead cells, FITC to identify $CD11c^{+}$ cells, PerCP/Cy5.5 to identify $CD80^{+}$ cells, APC to identify $CD86^{+}$ cells and eFluor™ 450 to identify MHC-$II^{+}$ cells, respectively.

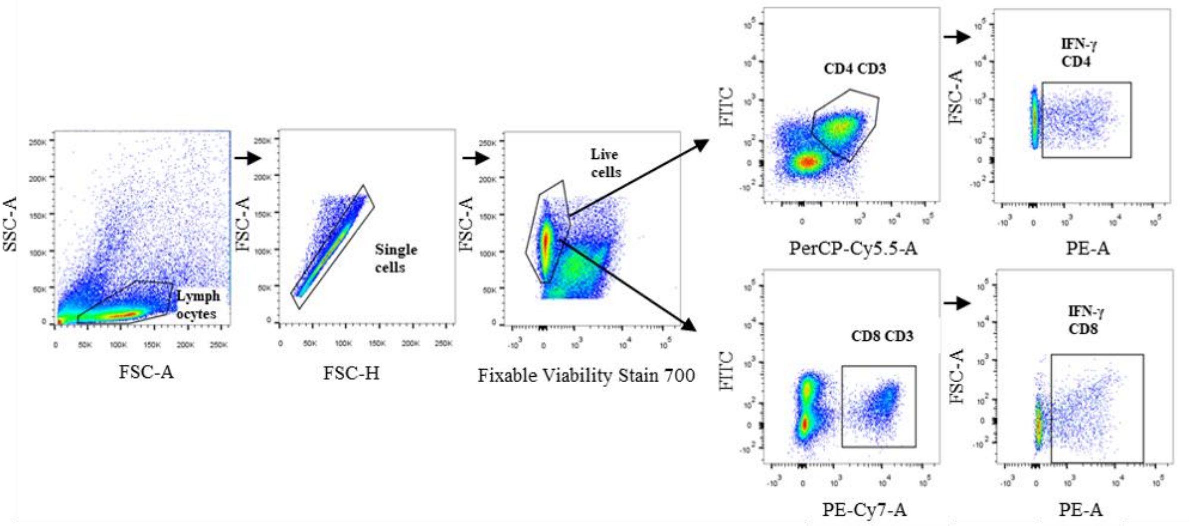


**Supplementary Fig. 10. Gating strategy for identification of interferon-γ (IFN-γ) secreting CD4$^+$ and CD8$^+$ T cells.** Initially, gating on cells based on FSC-A and SSC-A to exclude debris. Next, using FSC-A and FSC-H to exclude adhesive cells, Fixable Viability Stain 700 to exclude dead cells, APC/R700 to identify CD3$^+$ cells, PerCP/Cy5.5 to identify CD4$^+$ cells, PE-Cy7 to identify CD8$^+$ cells, and PE to identify IFN-γ$^+$ cells, respectively.

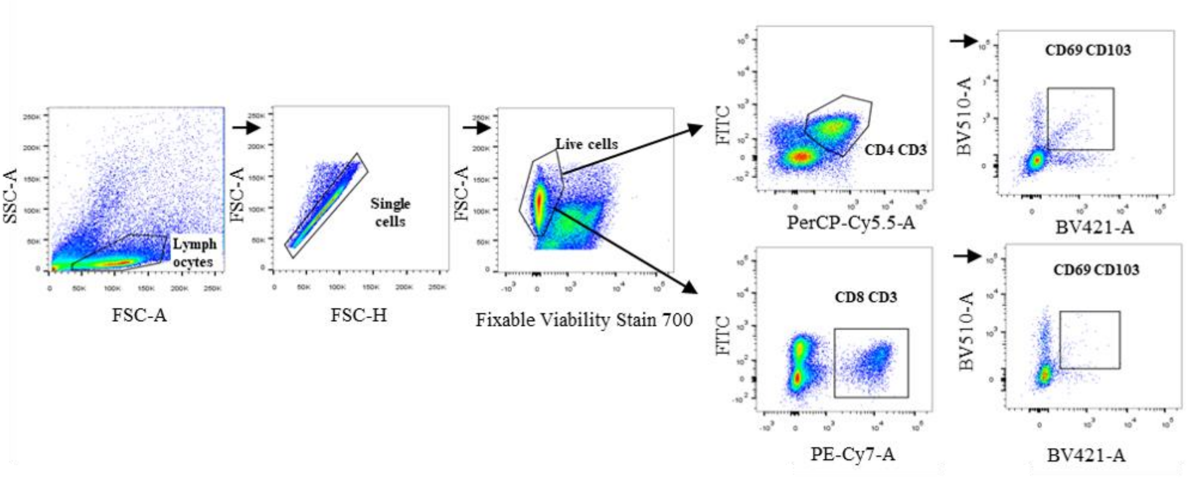


**Supplementary Fig. 11. Gating strategy for identification of Trm frequencies across CD4$^+$ and CD8$^+$ T cells.** Initially, gating on cells based on FSC-A and SSC-A to exclude debris. Next, using FSC-A and FSC-H to exclude adhesive cells, Fixable Viability Stain 700 to exclude dead cells, FITC to identify CD3$^+$ cells, PerCP/Cy5.5 to identify CD4$^+$ cells, PE-Cy7 to identify CD8$^+$ cells, BV421 to identify CD69$^+$ cells, and BV510 to identify CD103$^+$ cells, respectively.

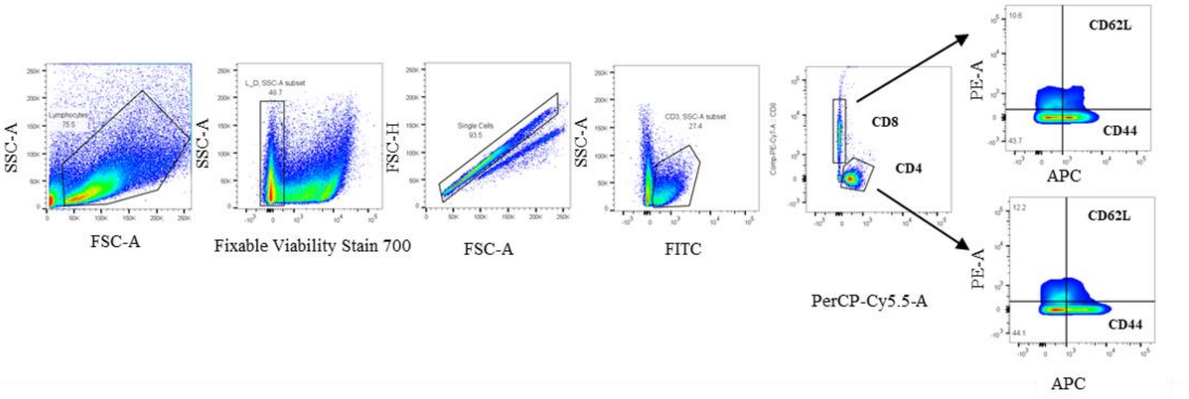


**Supplementary Fig. 12. Gating strategy for identification of Tcm and Tem across CD4$^+$ and CD8$^+$T cells.** Initially, gating on cells based on FSC-A and SSC-A to exclude debris. Next, using FSC-A and FSC-H to exclude adhesive cells, using Fixable Viability Stain 700 to exclude dead cells, using FITC to identify CD3$^+$ cells, PerCP/Cy5.5 to identify CD4$^+$ cells, PE-Cy7 to identify CD8$^+$ cells, APC to identify CD44$^+$ cells, and PE to identify CD62L$^+$ cells, respectively.
